# ‘The Thousand Polish Genomes Project’ - a national database of Polish variant allele frequencies

**DOI:** 10.1101/2021.07.07.451425

**Authors:** Elżbieta Kaja, Adrian Lejman, Dawid Sielski, Mateusz Sypniewski, Tomasz Gambin, Tomasz Suchocki, Mateusz Dawidziuk, Paweł Golik, Marzena Wojtaszewska, Maria Stępień, Joanna Szyda, Karolina Lisiak-Teodorczyk, Filip Wolbach, Daria Kołodziejska, Katarzyna Ferdyn, Alicja Woźna, Marcin Żytkiewicz, Anna Bodora-Troińska, Waldemar Elikowski, Zbigniew Król, Artur Zaczyński, Agnieszka Pawlak, Robert Gil, Waldemar Wierzba, Paula Dobosz, Katarzyna Zawadzka, Paweł Zawadzki, Paweł Sztromwasser

## Abstract

Although Slavic populations account for over 3.5% of world inhabitants, no centralized, open source reference database of genetic variation of any Slavic population exists to date. Such data are crucial for either biomedical research and genetic counseling and are essential for archeological and historical studies. Polish population, homogenous and sedentary in its nature but influenced by many migrations of the past, is unique and could serve as a good genetic reference for middle European Slavic nations.

The aim of the present study was to describe first results of analyses of a newly created national database of Polish genomic variant allele frequencies. Never before has any study on the whole genomes of Polish population been conducted on such a large number of individuals (1,079).

A wide spectrum of genomic variation was identified and genotyped, such as small and structural variants, runs of homozygosity, mitochondrial haplogroups and Mendelian inconsistencies. The allele frequencies were calculated for 943 unrelated individuals and released publicly as The Thousand Polish Genomes database. A precise detection and characterisation of rare variants enriched in the Polish population allowed to confirm the allele frequencies for known pathogenic variants in diseases, such as Smith-Lemli-Opitz syndrome (SLOS) or Nijmegen breakage syndrome (NBS). Additionally, the analysis of OMIM AR genes led to the identification of 22 genes with significantly different cumulative allele frequencies in the Polish (POL) vs European NFE population. We hope that The Thousand Polish Genomes database will contribute to the worldwide genomic data resources for researchers and clinicians.

## Introduction

An individual genome carries about four million single nucleotide variants and indels, and thousands of structural variants. These alterations in the DNA are mostly responsible for 0.1% of difference between genomes of two unrelated individuals (The 1000 Genomes Project Consortium 2015). Genetic variation is the reason for phenotypic differences - disease susceptibility, drug responses and physical traits such as height (Frazer et al. 2009). Mapping naturally occurring genetic variation is thus critical for studying human health and disease.

The completion of the Human Genome Project and subsequent large-scale projects, such as the HapMap (†The International HapMap Consortium 2003), the 1000 Genomes Project (The 1000 Genomes Project Consortium 2015), The ExAC Project (Exome Aggregation Consortium et al. 2016) and the TopMED (NHLBI Trans-Omics for Precision Medicine (TOPMed) Consortium et al. 2021) provided large datasets for human genomic variation studies. Most of these efforts were focused on the largest populations (https://sci-hub.se/https://www.nature.com/articles/nature18964) and dominated by participants of causasian ancestries, so despite sheer sizes, the information on the full spectrum of world’s genetic variation remains incomplete. With short read sequencing technology becoming cheaper and more accessible across the world, new large scale initiatives are launched, targeting thousands of genomes to fill in the gaps in our knowledge and build stronger foundations for precision medicine. In Europe, ‘The 1+ Million Genomes initiative’ (https://b1mg-project.eu/) is aiming to create the most comprehensive map of European diversity by bolstering up separate efforts of EU countries to sequence their populations. Similar projects are being held in the USA, ‘All of us’ project (https://allofus.nih.gov/) in Asia, GenomeAsia 100K Project (Wall et al. 2019) and Africa, 3MAG Project (ambitious Three Million African Genomes). Genetic variants derived from ‘healthy’ genomes on population scale are essential as a reference data for studying diseases, epidemiology, human diversity, evolution and migration. For example, the ClinVar (http://www.ncbi.nlm.nih.gov/clinvar/), Varsome (http://varsome.com) and HGMD (http://www.hgmd.org) databases store information about pathogenicity status of millions of small human variants (single nucleotide variants (SNVs) and indels), categorizing them to 5 groups according to ACMG guidelines (benign, likely benign, VUS, likely pathogenic and pathogenic). This clinical classification could not be created without population scale variant databases such as the 1000 Genomes or gnomAD (https://gnomad.broadinstitute.org). Similarly, Database of Genomic Variants (DGV) and DECIPHER aggregate the Structural Variants (SVs) and Copy Number Variants (CNVs) information for public use.

Here, we report the insights from the Thousand Polish Genomes (POL) project, an effort to sequence and characterize 1079 whole genomes of Polish individuals, including over 110 families. High quality and depth of the sequencing data (the average coverage above 35x) provide a unique repository of genetic variation in the Polish population. We characterized a wide spectrum of genetic alterations, including small and structural variants, as well as mitochondrial DNA. Taking into account that populations of central-European ancestry are underrepresented in the world’s large-scale genomic databases, the presented work complements international efforts to map human genetic variation. Additionally, an open allele frequency database^1^ will become an invaluable resource for clinical and population genetics, and biomedical research and demographic inference.

## Materials and methods

### Donors characteristics and sample collection

The studied population consisted of participants recruited within the “Search for genomic markers predicting the severity of the response to COVID-19” project related to genetic predisposition to COVID-19 severity. Samples were collected from 1079 individuals of Polish origin between April 2020 and April 2021. The whole cohort comprises three groups: (1) 441 participants from the Central Clinical Hospital of the Ministry of Interior and Administration in Warsaw. Majority of them (391 out of 441) were patients suffering from COVID-19 caused by SARS-CoV-2 coronavirus, peripheral blood samples from this group were collected during hospitalization. (2) 608 individuals were self-enrolled volunteers for the project. Samples from this group were collected in blood collecting facilities from all over Poland. Almost all Polish local administrative units (15 out of 16 voivodeships) were represented (Figure 1). (3) 30 individuals were collected in the Multidisciplinary Municipal Hospital of Józef Struś in Poznań from patients suffering from COVID-19. The study participants were unrelated individuals (N=943) or families. Among participating families (N= 117) there were 69 trios, 12 quartets, 5 pairs of siblings and 31 families consisting of one parent and a child.

**Figure 1.**
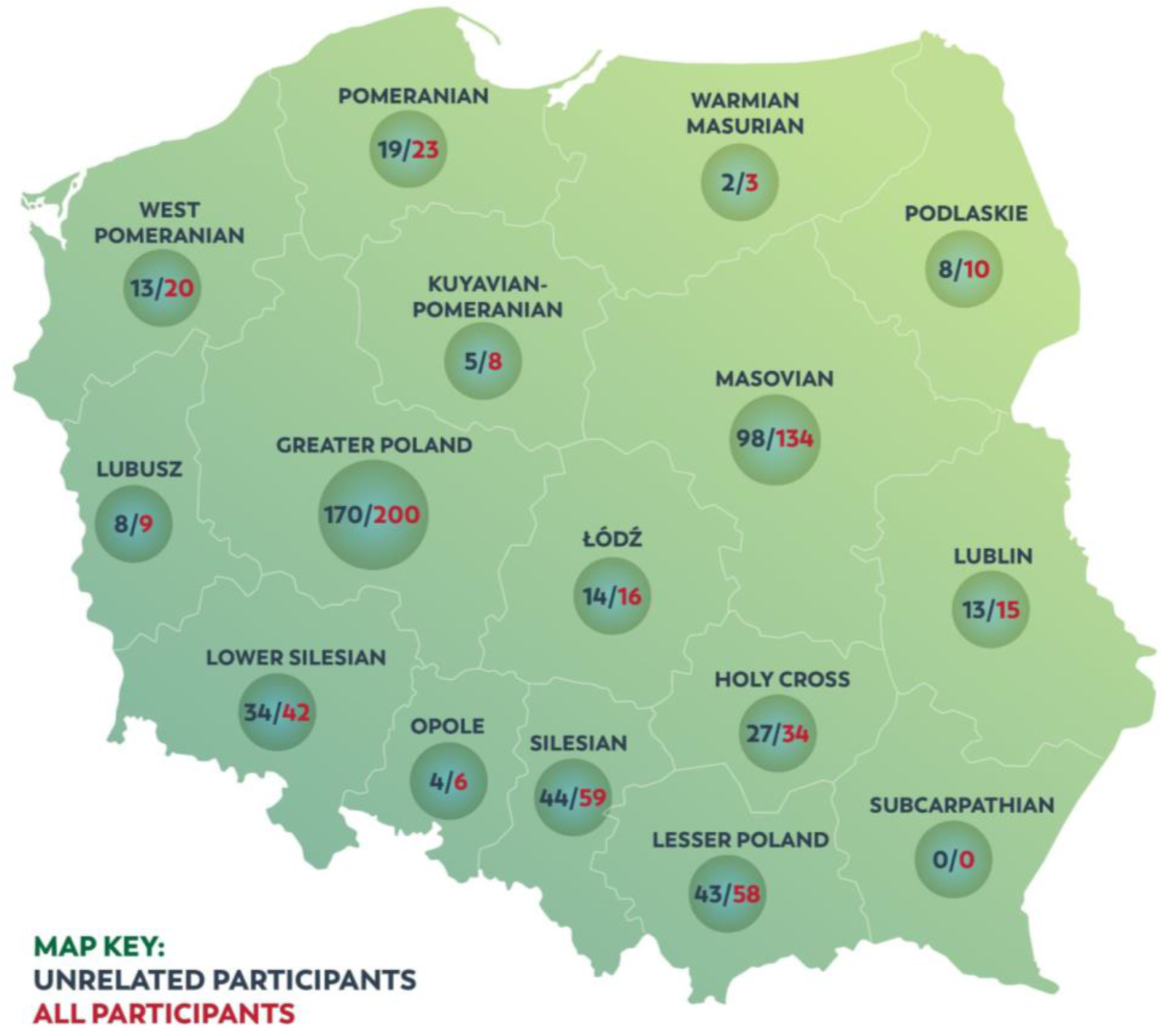
The map of 637 individuals enrolled in “The Thousand Polish Genomes Project” (related and unrelated ones) in respect of Polish voivodeships. It refers to individuals who volunteered to the project, from whom samples were collected in blood collection facilities from all over Poland.

From all participants of “The Thousand Polish Genomes Project” (POL) basic clinical history was collected in order to exclude genetically affected individuals. Data collected for all individuals included: gender, age, BMI, comorbidities (diabetes, hypertension, ischemic heart disease, stroke, heart failure, cancer, kidney failure/disease, hepatitis B, chronic obstructive pulmonary disease). Additional clinical data collected for 608 participants, not hospitalised for COVID-19 included: genetic disorders, flu vaccine, tuberculosis and measles vaccine, smoking habits, hepatitis C infection and others.

Peripheral blood samples from participants were collected in the EDTA-containing tubes. Whole process of the collection of biological material was monitored and performed in accordance with the protocol of blood collection, which is part of the Quality Management System.

### Compliance with Ethical Standards

All participants, or their guardians/parents (when the participants were under 18), provided their informed consent before collecting their blood samples and completed a clinical data form, which included a questionnaire about country of origin and chronic diseases. Only individuals without diagnosed severe genetic disorders were qualified for further studies. All procedures performed in studies involving human participants were in accordance with highest ethical standards, the Ethical approval for the study was obtained from Ethics Committee of the Central Clinical Hospital of the Ministry of Interior and Administration in Warsaw Ethics Committee (decision nr: 41/2020 from 03.04.2020 and 125/2020 from 1.07.2020). The study was in accordance with the 1964 Helsinki declaration and its later amendments and adhered to the highest data security standards of ISO 27001.The study is also compliant with the General Data Protection Regulation (GDPR) act.

#### Total Quality Management utilized in the study

The project was carried out in accordance with the Total Quality Management (TQM) methodology, which directly translates into the quality of results.

Before starting the project, the details were mapped - critical points were defined, such as reference ranges for collected biological material, its preparation, isolation, DNA concentration and quality, genomic sequencing (including quality control - QC). The legal and ethical transparency of the entire project was ensured, including confidentiality and data integrity as well as impartiality. Additionally, risk analysis was performed, possible difficulties were mapped and precise corrective actions have been planned.

## Methods

### Whole genome sequencing

High depth (>30x) PCR-free whole-genome sequencing was applied to all samples. Genomic DNA was isolated from the peripheral blood leukocytes using a QIAamp DNA Blood Mini Kit, DNeasy Blood & Tissue Kit (Qiagen), Blood/Cell DNA Mini Kit (Syngen) and Xpure Blood Kit (A&A Biotechnology) according to the manufacturer’s protocols. The concentration and purity of isolated DNA was measured using the NanoDrop™ spectrophotometer, the DNA integrity was evaluated with electrophoresis. The sequencing library was prepared by Macrogen Inc. (Seul, South Korea) using TruSeq DNA PCR-free kit (Illumina Inc., San Diego, California, United States) and 550 bp inserts. Quality of DNA libraries were measured using 2100 Bioanalyzer from Agilent Technologies. Subsequently Whole Genome Sequencing (WGS) was performed on the Illumina NovaSeq 6000 platform using 150 bp paired-end reads, with median average depth of 35.72X (Supplementary Table S1).

### WGS data processing

Quality of the sequenced reads was confirmed using FastQC v0.11.7^2^ and the reads were subsequently mapped to the GRCh38 human reference genome using Speedseq framework v.0.1.2 (Chiang et al. 2015), encompassing BWA MEM 0.7.10 (H. Li 2013) mapping, SAMBLASTER v0.1.22 (Faust and Hall 2014) duplicate removal, and Sambamba v0.5.9 (Tarasov et al. 2015) sorting and indexing. Mapping coverage was calculated using Mosdepth 0.2.4 (Pedersen and Quinlan 2017) and included reads with MQ>0. Single nucleotide variants and short indels in the nuclear genome were detected using DeepVariant 0.8.0 (Poplin et al. 2018) and jointly genotyped with GLnexus (Yun et al. 2021). Next, multiallelic variants were decomposed and normalized using BCFtools (Danecek et al. 2021). Small variants in the mitochondrial genome were called using Freebayes v.0.9.21 (Garrison and Marth 2012), and mitochondrial haplogroups inferred using Haplogrep2 software (Weissensteiner et al. 2016). Structural variants were called and jointly genotyped with Smoove v.0.2.6^3^.

All variants were annotated using Ensembl Variant Effect Predictor v.97 (McLaren et al. 2016), including references to databases of genomic variants from ClinVar v. 201904 (Landrum et al. 2018) and dbSNP build 151, variant population frequencies from the 1000 Genomes Project (Auton et al. 2015), GnomAD v2.0.1 and v3.0 (Karczewski et al. 2020), and pathogenicity scores including Polyphen-2 (Adzhubei et al. 2010), SIFT (Sim et al. 2012), and DANN (Quang, Chen, and Xie 2015). All WGS data processing was automated with a Ruffus pipeline framework (Goodstadt 2010), and some analysis steps were parallelized with GNU parallel tool (Tange 2011).

#### Variant analysis

Small variants of 943 unrelated individuals were extracted from the normalized multisample VCF file using BCFtools (Danecek et al. 2021). Sites with genotyping rate below 90% and invariant in the group were excluded. The resulting dataset, with average call rate per individual of 99%, was used in all subsequent analyses. Joint called structural variants were also filtered for related individuals, and subsequently annotated and filtered according to Smoove^4^ authors’ recommendations, i.e. heterozygous calls with MSHQ>=3, deletions with DHFFC>=0.7, and duplication calls with DHFFC<=1.25 were excluded from the analysis. We also required at least one split-read per allele in the cohort. Variant statistics were generated with BCFtools (Danecek et al. 2021), and custom scripts. Data processing and visualization was programmed in R (R Core Team 2020) with the tidyverse package (Wickham et al. 2019). Small and structural variant sets along with allele frequencies were released as a publicly available resource - The Thousand Polish Genomes database^5^.

#### Mendelian inconsistencies

Small variants violating Mendelian inheritance pattern were identified in 93 child-parents trios, and subsequently filtered using BCFtools (Danecek et al. 2021). For *de novo* candidates we selected sites covered by at least 10 reads in all family members, with minimum 5 alternative allele reads in the child and 0 such reads in the parents, and with minimum alt-allele depth ratio of 0.25.

#### Runs of homozygosity

Runs of homozygosity were identified using a hidden Markov model approach (Narasimhan et al. 2016) implemented in BCFtools (Danecek et al. 2021). We analysed RoH Identified on autosomal chromosomes, requiring more than 50 markers per RoH, and average marker quality above 25.

#### Recessive carriers identification

The list of Autosomal Recessive disease-associated genes was obtained from the OMIM database^6^. Variants located within these genes were extracted from the POL and gnomAD v3.0 cohorts and annotated using VEP (release 101), including ClinVar annotation (release date 17-05-2021). Of those, we selected rare (gnomAD_AF < 0.01%), nonsynonymous (VEP IMPACT = ‘HIGH’ or ‘MODERATE’), pathogenic or likely pathogenic (according to ClinVar) variants. For each gene, we calculated cumulative allele frequencies of previously selected variants in POL and gnomAD NFE cohorts. To test whether the cumulative frequencies in particular genes differ between both cohorts we used chi-square test and applied Bonferroni correction for multiple testing to get adjusted P-values.

We downloaded gene panels data from panelapp website^7^ and selected high confidence gene-disease associations (annotated as “Expert Review Green”) with ModeOfInheritance defined as “BIALLELIC”. For each panel we calculated the sum of cumulative allele frequencies of pathogenic/likely pathogenic variants (obtained in the previous step) in genes included in the panel. Similarly to the gene level analysis we tested the difference between POL and gnomAD NFE populations using a chi-square test followed by Bonefroni correction.

#### Population analysis

We evaluated the homogeneity of the Polish population (POL) using three different approaches. In the first step we checked whether all the analyzed POL samples belong to the European population. We used common variants occurring in the 1000G project and our cohort data and further pruned them to Minor Allele Frequency (MAF) over 1% and to be in linkage equilibrium. In the second step, we performed Principal Components Analysis (PCA) on a 1000G dataset and projected them into POL genotype data. We trained a random forest model with the first six principal components (PCs) on the 1000G dataset and predicted the ancestry of POL individuals. In the third step, in order to find out which European population is the closest to POL we used the subpopulations information available in 1000G data set (Utah Residents (CEPH) with Northern and Western European ancestry [CEU], Finnish in Finland [FIN], British in England and Scotland [GBR], Iberian population in Spain [IBS], Toscani in Italy [TSI]) and repeated the PCA and prediction based on random forest analyses as was described above. The only exception was to use the first two PCs in training and prediction of the random forest.

We also performed population structure analysis using ADMIXTURE (Alexander, Novembre, and Lange 2009) with the unrelated individuals from our cohort, filtered and pruned as described above, with the populations from the 1000G dataset, as well as with the five European populations. The analysis was performed in the unsupervised mode for K ranging from 2 to 7, and cross-validation error was used to determine the best value for K.

To calculate the average fixation index between two groups of individuals from SNPs we used Fst statistics calculated using the formula proposed by Weir and Cockerham (Weir and Cockerham 1984). The calculations were performed in Plink software (Purcell et al. 2007). We also compared the Polish population with all populations available within the 1000 Genomes Project and checked whether Fst patterns were consistent with the continental and the populational clustering and with previously obtained estimates. We also calculated an average Fst statistics over each sliding window built from 1,000 SNPs. We used the two first PCs and the Density-Based Spatial Method implemented in Python to determine potential subpopulations within the Polish population.

## Results

### Characteristics of the POL cohort

Median age of participants was 45 (2-98), with slight predominance of males (618 vs 461). The analysis of clinical data showed that the most common chronic diseases reported by participants of the POL cohort were: hypertension (13%), cancer (4,6%), diabetes (4%) and hypothyroidism or Hashimoto’s disease (3%). No health problems (excluding COVID-19 infection which was a major focus of patients recruitment) were reported by 86% of participants. The whole cohort consists of **1079** individuals of Polish origin, **943** of which are unrelated. The figure below (Fig. 2) shows the age distribution of all unrelated participants.

**Figure 2:**
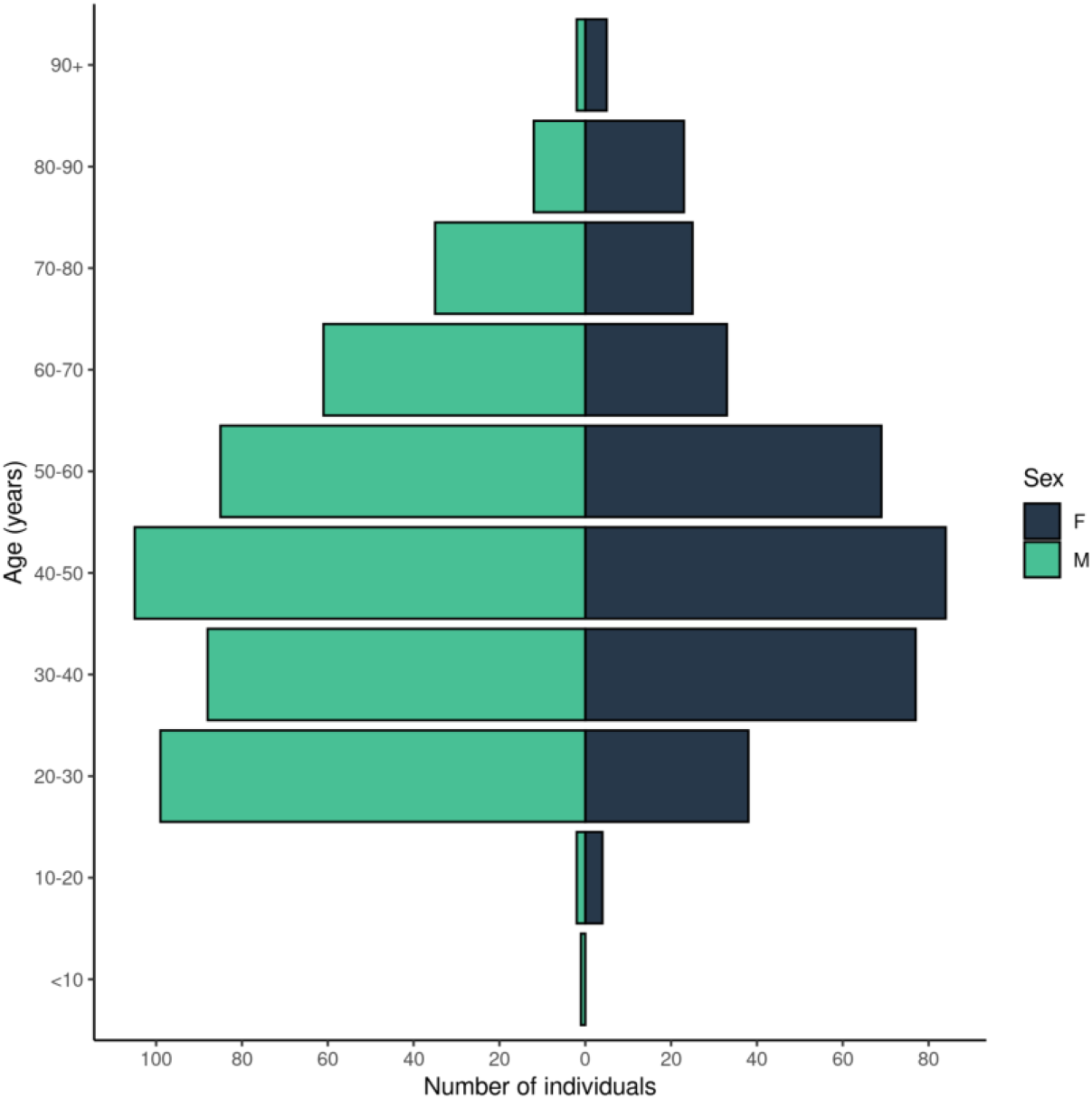
Age distribution of unrelated study participants.

### Genetic variation in the Polish population

We processed sequencing data from 1079 individuals, totalling 878.4 billion read-pairs corresponding to an average 35.72x mean coverage per genome (Supplementary Fig. S1; Supplementary Table S1). In all samples 92% of the genome was covered with at least 10 reads, and in 41% of samples 90% of the genome was covered by at least 20 reads. We detected and genotyped small and structural variants, runs of homozygosity, mitochondrial haplogroups, and Mendelian inconsistencies, all of which are described in detail below. Except for Mendelian inconsistencies, in all analyses we included data for 943 unrelated individuals, a group referred to as the POL cohort below. Allele frequencies of small and structural variants from this cohort constitute The Thousand Polish Genomes database^8^, that we release openly for academic research.

### Small and structural variants

We identified a group of small variants including a total of 31.24M single nucleotide variants (SNV) and 5.63M small insertions and deletions. An individual genome contained on average 3.71M (3.63 - 3.77) single nucleotide substitutions of which 2.22M (2.08 - 2.36) were heterozygous and 1.49M (1.38 - 1.55) homozygous. Together with, on average, 0.77M (0.75 - 0.78) short insertions and deletions, we found 4.48M small variants per individual, of which 16,473 were private variants (Supplementary Table S2).

An average individual genome also carried 2,602 large deletions (2,431-2,724), 106 duplications (86-128), and 76 inversions (56-91). The Thousand Polsih Genomes database holds a total of 59,823 structural variants, of which 21,492 were marked as high quality (18,971 deletions, 2,148 duplications, and 373 inversions; Supplementary Table S3 and S4). Majority of deletions were shorter than 1kb (63.6%; 93.1% were <10kb), with peaks at 50bp and 300bp (Supplementary Fig. S2 A-C). Distribution of duplication sizes showed two peaks, at 200bp and at 10kb. Only 10 duplications (0.5%) exceeded 1Mb in length. Inversions, being the most rare category of the three structural variant types, displayed a clear length peak below 1kb, and the highest fraction of events longer than 1Mb (2.7%; N=10). Median length of the genomic sequence affected by SVs in a single individual was 12.6Mb (2.94Mb, 2.34Mb, and 7.34Mb for deletions, duplications, and inversions respectively), which exceeds the total DNA content affected by SNVs.

### Distribution of allele-frequencies

We compared distribution of allele frequencies among different variant types: substitutions, short indels, and structural variants. The results presented in Fig. 3 indicate that the majority of substitutions (50.8%) were private variants, observed in a single individual (MAF ∼0.05%). For deletions and duplications singletons constituted 41 and 43.7% respectively, while for the inversions only 26%. Common (MAF>1%) structural variants in our cohort were most abundant among inversions (50%), and least abundant among duplications (15.6%). High frequency small variants fell in the middle with 29.4% common for SNVs and 38.7% for indels. Interestingly, fixed alleles (allelic frequency equal to 100%) were present among all three types of structural variants (0.1-0.5% of all SVs) and completely absent among small variants. A summary of small variant counts with breakdown into three allele frequency tiers is shown in Table 1.

**Table 1:** Summary of variant counts in three population allele frequencies.

| VARIANT_CLASS | AF | IMPACT |  |  |  |
| --- | --- | --- | --- | --- | --- |
|  |  | HIGH | LOW | MODERATE | MODIFIER |
| deletion | >0.5% | 500 | 1 090 | 680 | 1 697 005 |
| insertion |  | 327 | 1 280 | 630 | 1 934 758 |
| SNV |  | 1 446 | 45 903 | 39 096 | 15 294 308 |
| deletion | 0.1-0.5% | 813 | 582 | 859 | 885 811 |
| insertion |  | 407 | 648 | 573 | 1 010 671 |
| SNV |  | 1 723 | 32 131 | 41 239 | 9 509 163 |
| deletion | <0.1% | 2 600 | 946 | 1 684 | 1 383 954 |
| insertion |  | 1 219 | 773 | 983 | 1 062 161 |
| SNV |  | 4 967 | 68 293 | 100 838 | 19 225 909 |

**Figure 3:**
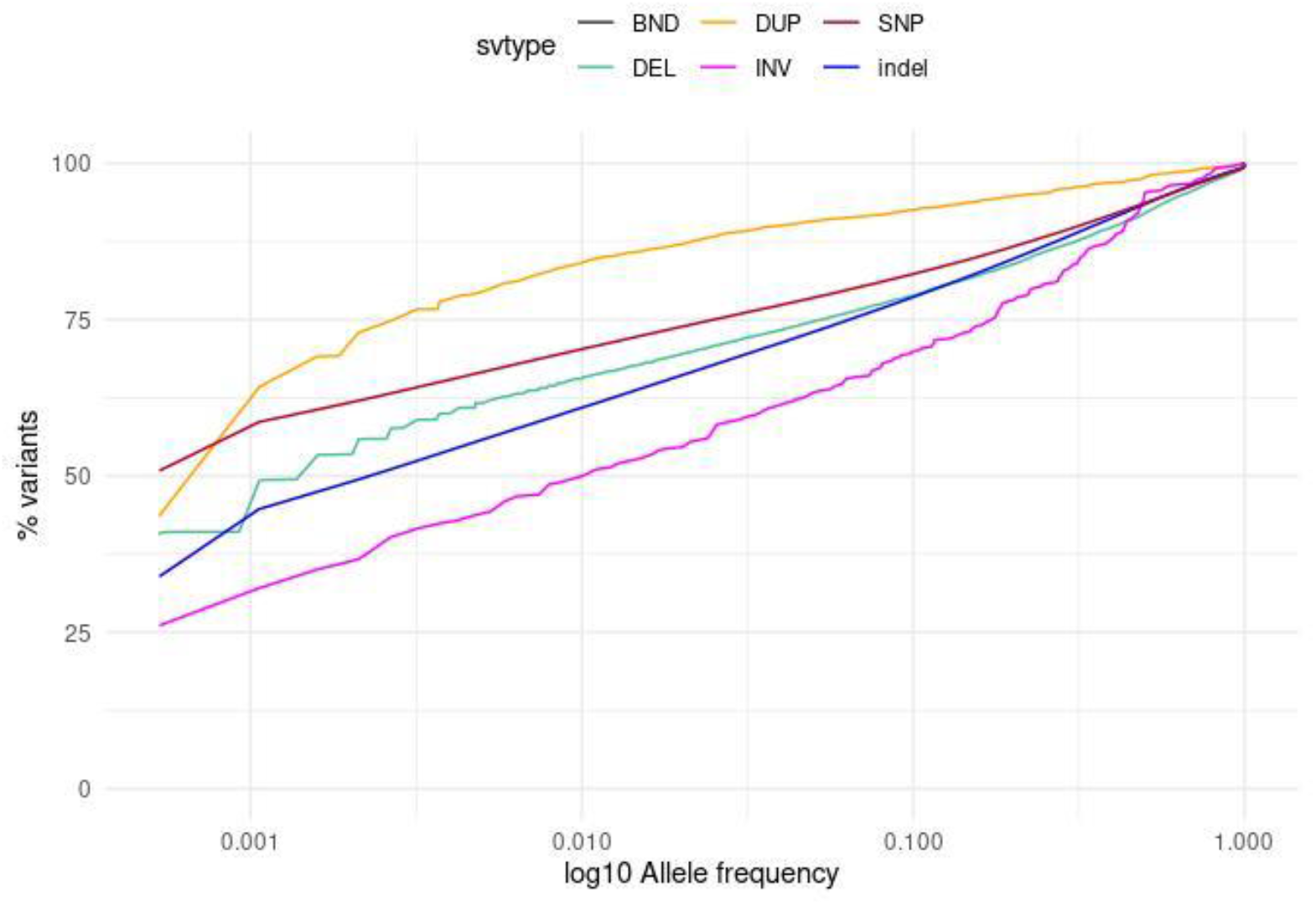
Distribution of allele frequencies across different variant types.

### *Denovo* variation in family-based analysis

Using small variant genotypes in 93 parents-child trios we identified on average 11,348 (10,128-13,224) Mendelian inconsistencies per trio. Considering only variants with MAF<1% in the entire cohort, a child carried on average 347 substitutions (284-456), and 1,127 indels (977-1,376). A total of 194 rare *denovo* SNVs affected the protein coding sequence (114 missense, 53 synonymous, 23 splice-site or region, and 4 stop-gain) corresponding to 0-4 exonic SNVs in 89 trios, and more than 4 in four trios (5-8). On average, a child carried 2,1 *denovo* SNVs with MAF<1%.

### Runs of homozygosity

On average, an individual genome contains 1.1 runs of homozygosity (RoH) longer than 100kb, spanning a total of 284Mb. RoH were uniformly distributed along the chromosomes, with the exception of centromeric and near-centromeric regions (Fig. 5). Mean size of a RoH was 252.7kb (100kb-14Mb), and counts and average sizes of RoH on particular chromosomes can be found in Supplementary F ig. S3 and Supplementary Fig. S4. The steepest growth in cumulative length of RoHs (Fig. 4) was observed between 20kb and 200kb, and RoH longer than 200kb represented less than a quarter of the homozygous sequence (median 88Mb).

**Figure 4:**
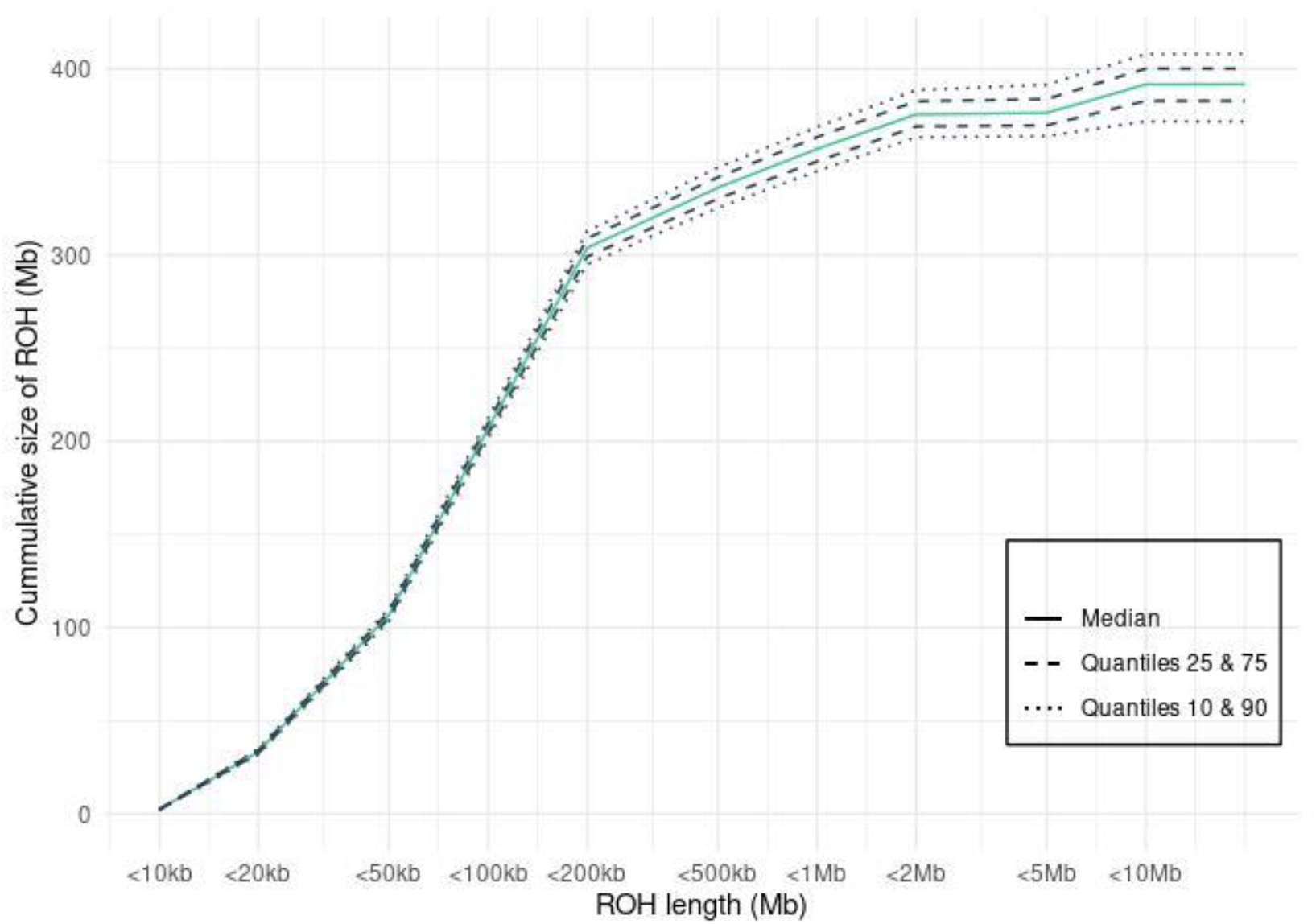
Cumulative length of RoHs.

**Figure 5:**
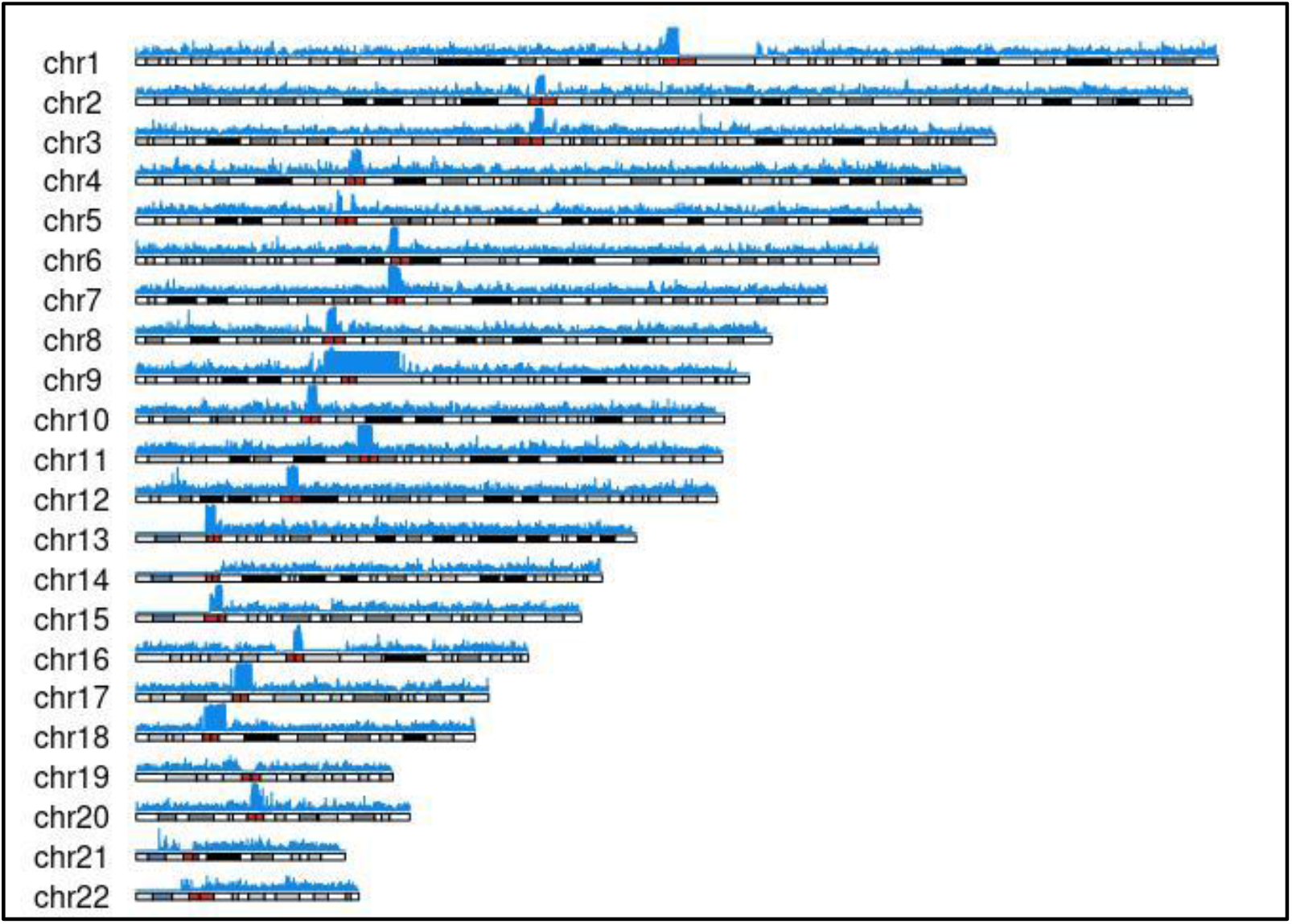
Chromosomes coverage by runs of homozygozity in the POL cohort.

### Mitochondrial haplogroups

Using variant calls in the mitochondrial genome, we inferred haplogroups among the unrelated individuals. In 930 individuals with high quality haplogroup assignment the most abundant haplogroup was H with 410 (44.1%) representatives, U with 161 (17.3%), J with 92 (9.9%), and T with 83 (8.9%) individuals (Fig. 6). The largest H sub-haplogroup was H1 (N=128; 31.2% of the H haplogroup), and a similar number of individuals was divided between subclades H2, H5, H6 and H11 (N=116; together 28.3% of the H haplogroup). The second most abundant sub-haplogroup in the cohort was U5 with 98 (10.5%) individuals.

**Figure 6:**
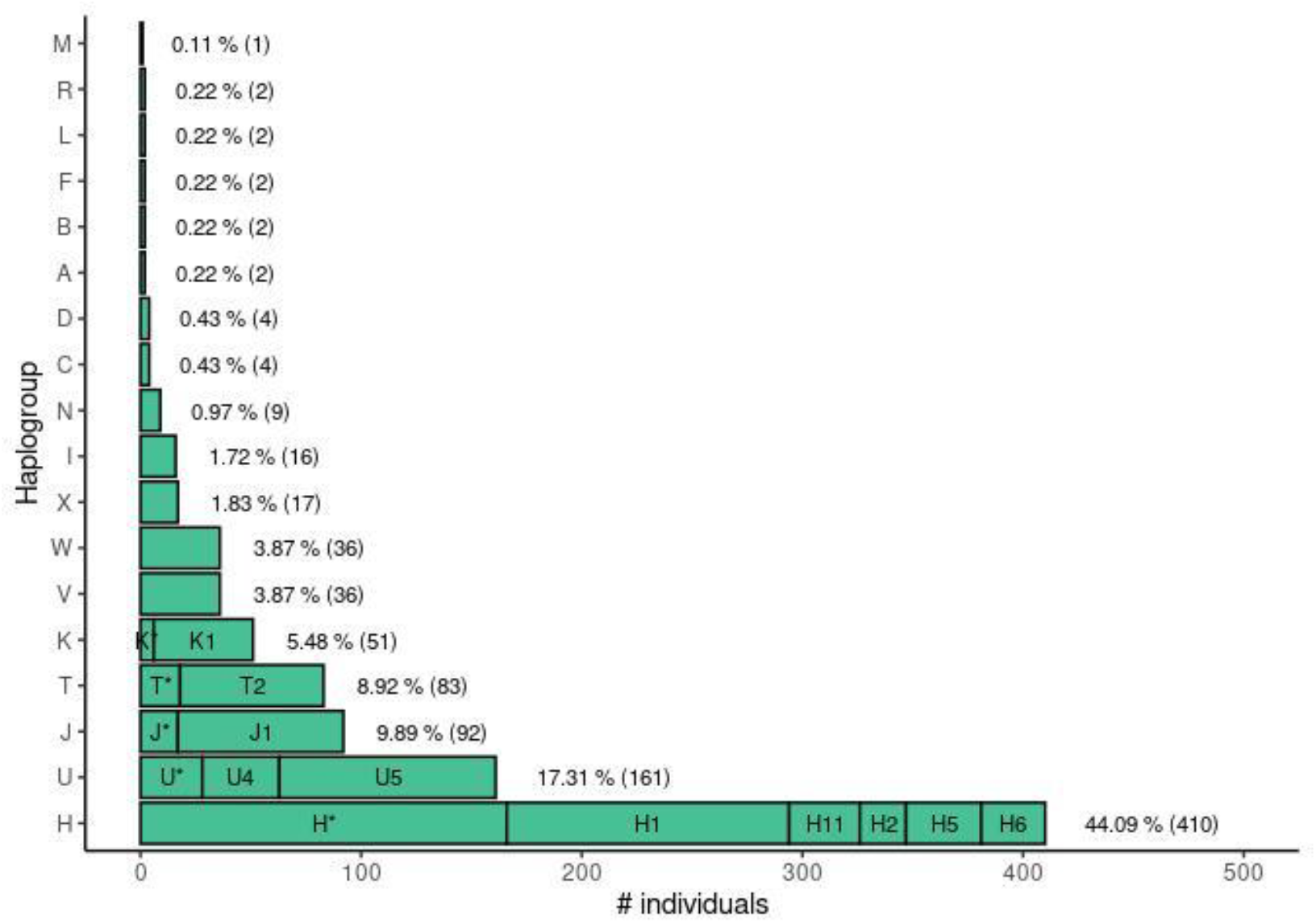
Haplogroups distribution in POL.

### Functional impact of the variants

#### Impact of small variants on protein function

Depending on the impact on the genomic sequence, variants were distributed non uniformly across the allele-frequency spectrum. Variants in protein coding regions were depleted in higher frequencies (Supplementary Fig. S5) compared to non-coding variation (e.g. intergenic, intronic). Larger differences were observed among exonic variants (Fig. 7), where protein altering variation (nonsense variants, frameshift indels, and missense variants) was associated with lower allele frequencies in contrast to UTR, synonymous, and non-exonic variants. A summary of variant counts, SNVs and indels, divided into three population frequency tiers and categorized by functional impact is presented in Table 1.

**Figure 7.**
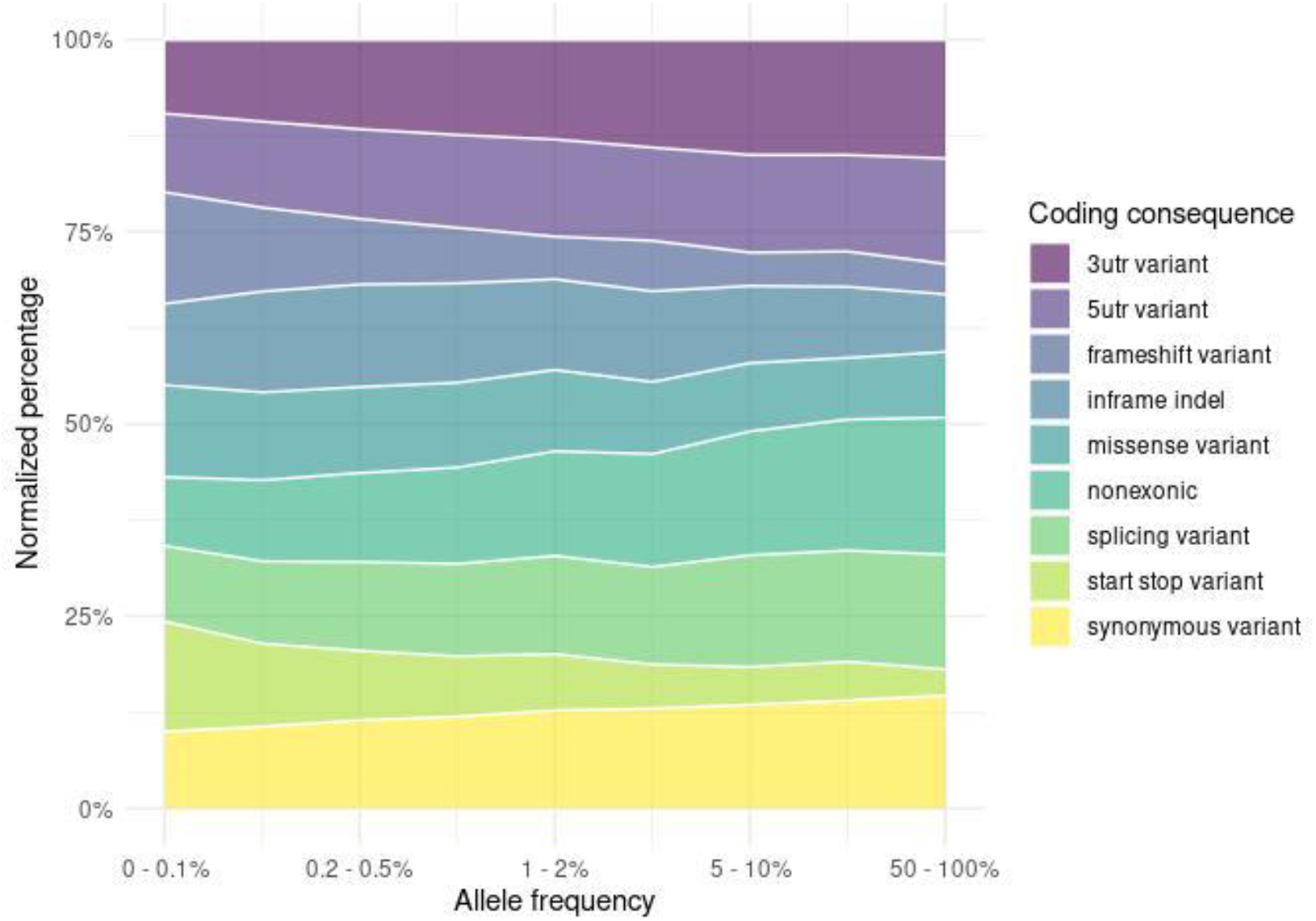
Distribution of exonic variant consequences across allele frequency spectrum.

#### Impact of rare variants on diseases

In order to infer the burden of disease causing variants in the Polish population, we considered three rare variant sets: (1) variants annotated in the ClinVar database as pathogenic or likely pathogenic; (2) high-impact variants in genes from the ACMG gene list; (3) variants associated with autosomal recessive (AR) diseases according to the OMIM database^9^. The details of the three sets and the lists of corresponding genes are described below.

In the first approach, we identified 736 rare variants (MAF<0.1% in gnomAD v3.1) reported in ClinVar as pathogenic or likely pathogenic. Selected variants were divided into four groups by the review status, which is the assertion of clinical significance: (1) 282 variants with a low confidence level (provided by a single submitter, interpreted with assertion criteria and evidence or provided by multiple submitters, but with conflicting interpretations of pathogenicity); (2) 432 variants with higher clinical significance (submitted to ClinVar by two or more submitters with the same interpretation), for example rs11555217 in *DHCR7* gene, described in more details below; (3) 20 pathogenic variants with high clinical significance (variants reviewed by an expert panel) and (4) 2 variants with the highest level of significance (assessed as “practice guideline” assertions). 533 (72.4%) of these variants were observed in a single person and 203 variants were carried by 2-22 individuals (for instance variants in *PAH, OTOG* and *CBS* gene).

In the second approach, we evaluated clinically relevant ‘incidental findings’ variants according to novel American College of Medical Genetics and Genomics (ACMG) recommendations. The most recent version of ACMG (v.3.0) guidelines was published in May 2021 (ACMG Secondary Findings Working Group et al. 2021). Using this list we identified 50 variants in POL having a high impact on genes. After manual curation by certified medical genetic scientists, 13 variants were assessed as pathogenic and 3 as likely pathogenic, affecting 18 of 943 (1.9%) unrelated healthy individuals in our cohort. 15 (94%) of these variants were observed in a single person and were mostly related to cancer predisposition genes such as *MUTYH, MSH6, BRCA2*, and *PALB2*. In one case the variant was carried by three individuals (rs1217805587 in *BRCA1* gene). Frequencies of these variants did not differ significantly from the gnomAD NFE population.

In the third approach, we extracted 2 512 genes associated with Autosomal Recessive (AR) diseases and then gene panels containing AR disease genes from the OMIM database. We found 26 288 and 73 036 rare (AF < 0.01 % in GnomAD), nonsynonymous (IMPACT= ‘HIGH’ or ‘MODERATE’) variants in the POL and GnomAD cohorts respectively. Of those, 705 (POL) and 10 116 (gnomAD) variants have been annotated in ClinVar database as pathogenic or likely pathogenic. Per-gene analysis of the cumulative allele frequencies of these variants revealed the existence of significant differences (Chi-square test, P value < 0.01) in 22 genes between POL and gnomAD NFE populations (Supplementary Table S5, Figure 8).

**Figure 8:**
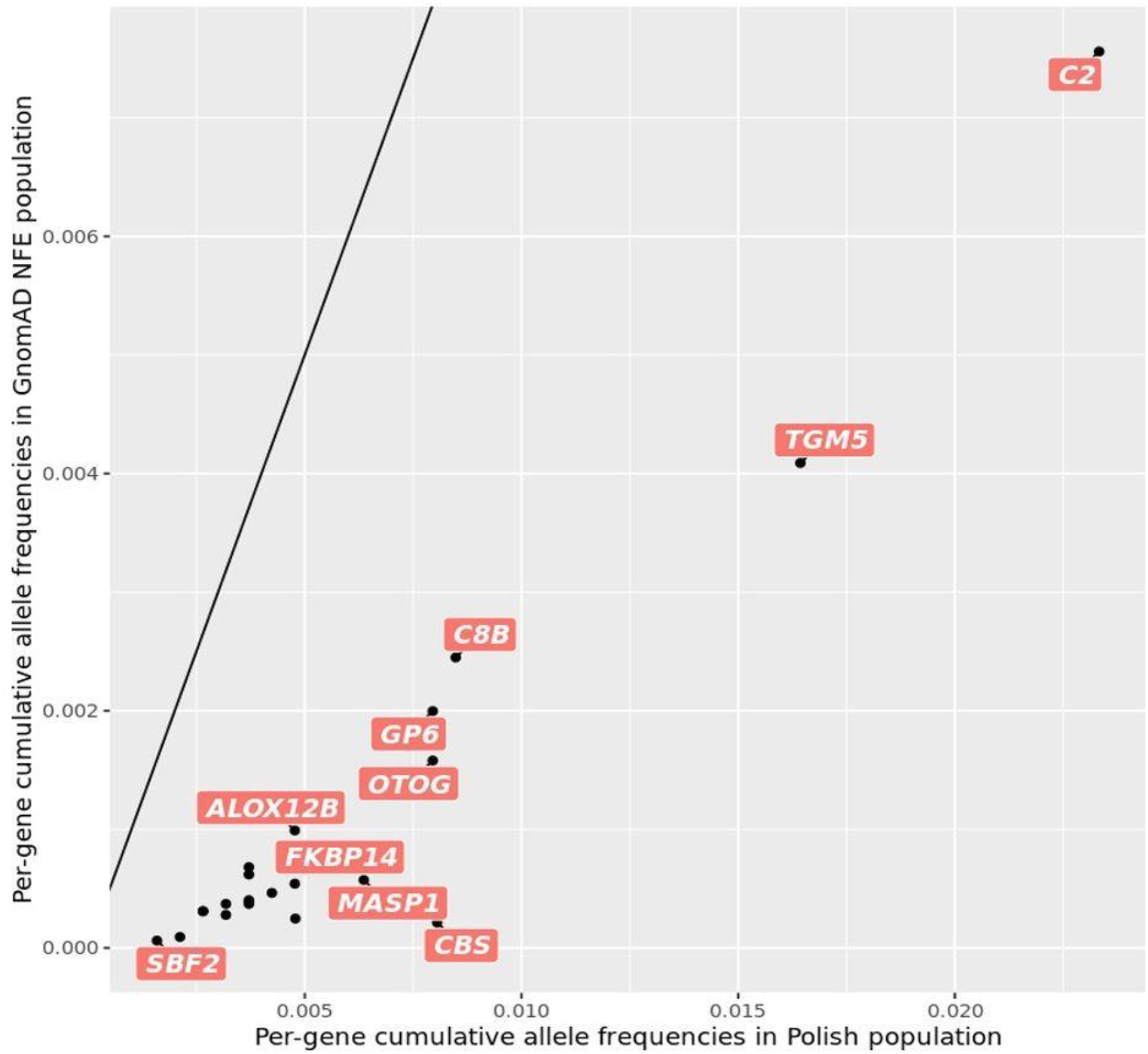
Cumulative allele frequencies of selected ClinVar variants in 22 AR genes.

A gene with one of the the highest log2 fold change of over 3.3 in the POL cohort was *XPA*, encoding a protein that plays a crucial role in the nucleotide excision repair process, a specialized type of DNA repair (Musich, Li, and Zou 2017). Pathogenic variants in this gene cause Xeroderma Pigmentosum (XP) type A, one of seven complementation groups of this disorder. There is a variability in prevalence within complementation groups with *XPC* being the most common complement type in the United States, Europe, and North Africa while *XPA* is the most common type in Japan (Black 2016). Our results suggest that *XPA* pathogenic variants are more common in the Polish population vs gnomAD NFE populations.

Other genes with pathogenic variants that were overrepresented in the Polish populations include: *C2, TGM5, C8B, CBS, OTOG, GP6, MASP1, C19orf12, SORD, ALOX12B, FKBP14, PIGV, ABCB11, ENAM, PROP1, HPGD, SLC26A3, SLC19A1, AP4B1, STAG3*, and *SBF2*.

Next, we analyzed 248 gene panels from panelApp, containing 1 823 AR disease genes. We identified significant differences (Chi-square test, P-value < 0.01) in cumulative allele frequencies between POL and NFE populations in 30 gene panels (Fig. 9; Supplementary table S6). Among these, “Optic neuropathy” panel was the most enriched in pathogenic variants in POL in comparison to the NFE population.

**Figure 9.**
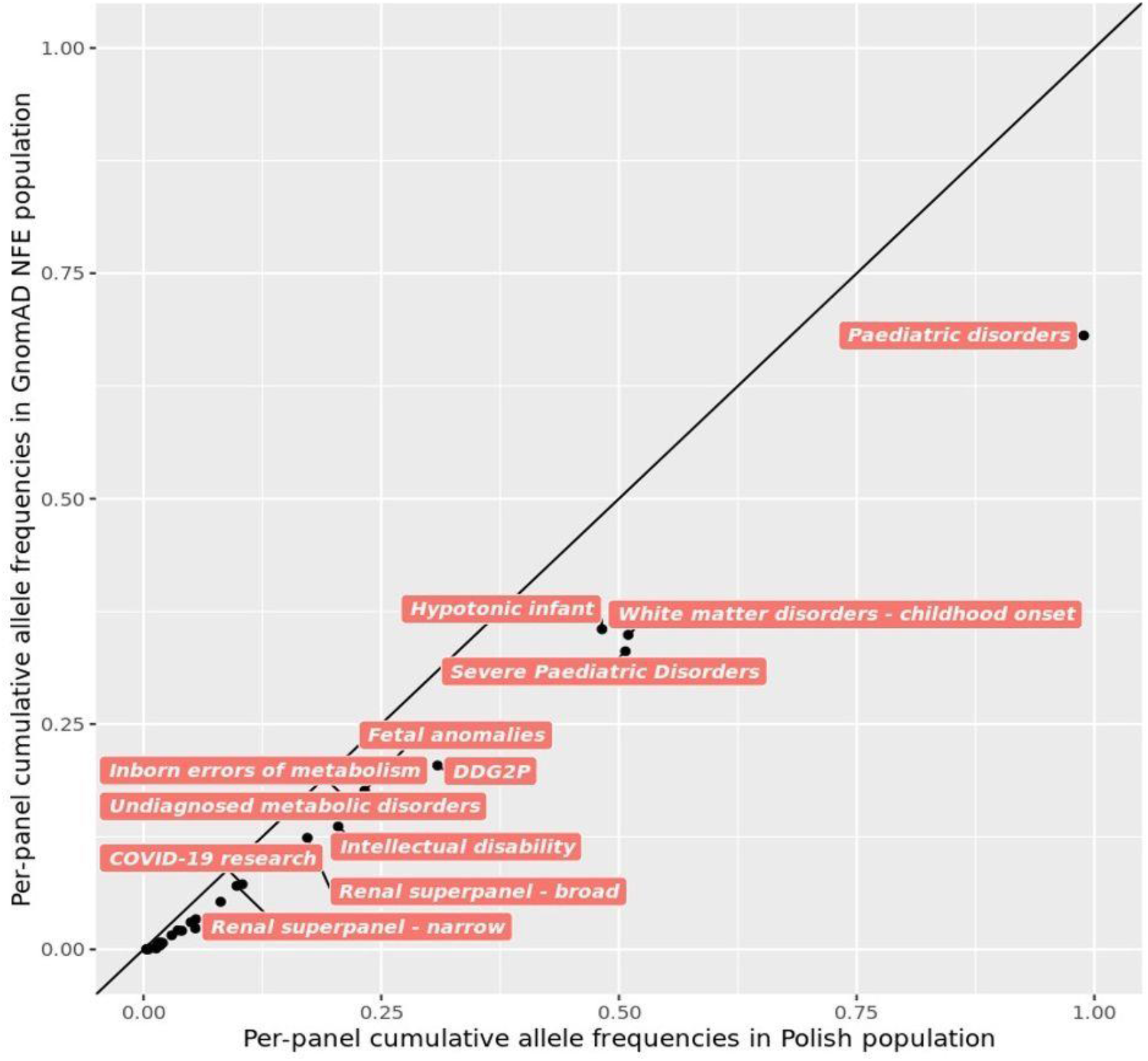
Cumulative allele frequencies of selected ClinVar variants in AR gene panels.

#### Selected disease causing variants

In our study, we took a closer look at variants characteristic for the Eastern European or the Slavic populations. We selected three well known diseases, as examples of POL database utility for verification of the genetics of well studied severe diseases.

The most frequent mutation from the list of pathogenic or likely pathogenic variants reported in ClinVar was found in 22 unrelated participants of POL (MAF=1.17%). It was the NG_012655.2:g.12031G>A, p.Trp151Ter variant (rs11555217) located in the *DHCR7* gene which mutations are responsible for Smith-Lemli-Opitz syndrome (SLOS). It is an autosomal recessive disorder characterized by failure to thrive, microcephaly, intellectual disability, cleft palate and multiple birth defects, such as syndactyly of second and third toes, dysmorphic facial features or heart defects (Ryan et al. 1998). Global MAF for the p.Trp151Ter variant is 0.07% (GnomAD v3.1.1) which is significantly lower than the frequency observed in our cohort (Fig. 10A). SLOS occurs as a common recessive disorder in Europe, but is almost absent in most other populations. Previous studies showed that mutations causing this disorder differ among European populations. The founder p.Trp151Ter variant was the most frequent mutation in Polish SLOS patients compared to patients from other subpopulations such as German or British (Witsch-Baumgartner et al. 2001).

**Figure 10.**
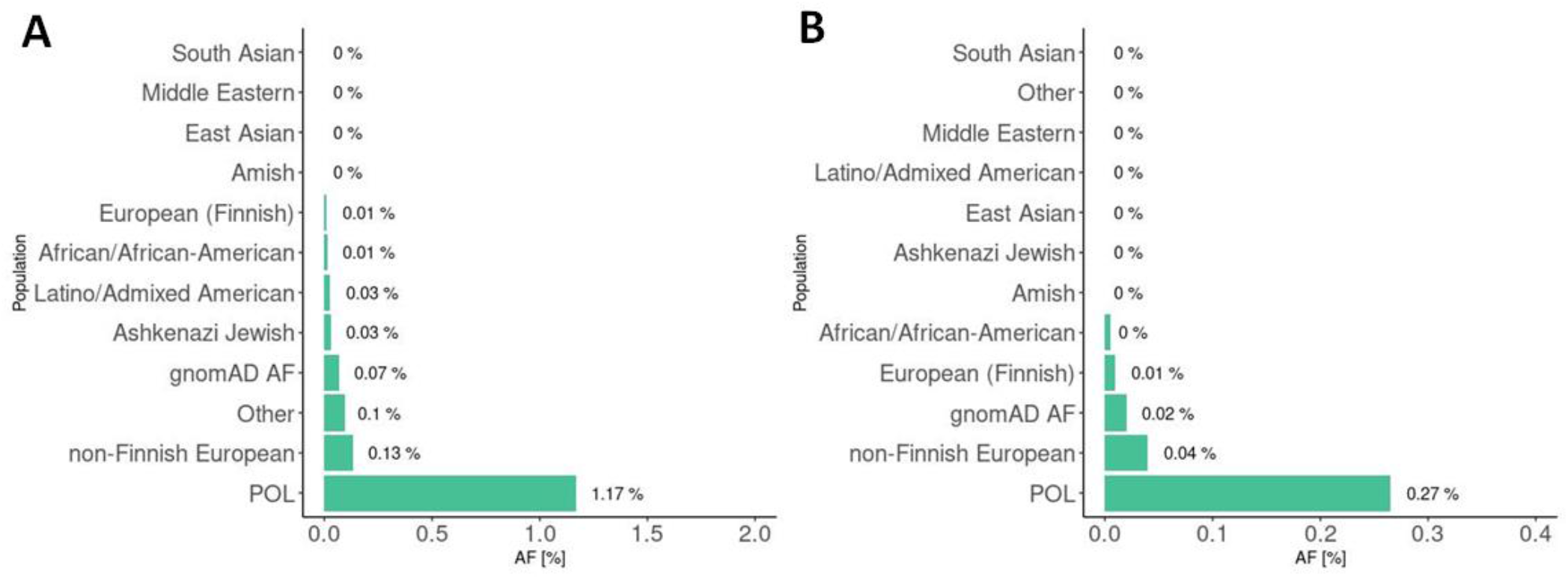
Frequency of selected variants in Polish (POL) and gnomAD v.3.1.1 populations: the p.Trp151Ter variant (rs11555217) located in the *DHCR7* gene (A) and the 657del5 variant located in the *NBN* gene (B).

The second interesting example identified in our study is the rare, recessive NM_002485.5:c.657_661del, resulting in p.Lys219fs variant (rs587776650) located in the *NBN* gene which mutations are responsible for Nijmegen breakage syndrome (NBS). NBS is characterized by microcephaly, immunodeficiency and very high predisposition to lymphoid malignancy. It was reported that NBS heterozygosity is also responsible for increased incidence of tumors, especially lymphoma (Seemanová 1990), meaning that the NBS variant contributes to the cancer frequency in a given population. The disease occurs worldwide, but its prevalence was reported to be significantly higher in Poland, Ukraine, Czech Republic and Russia (Kostyuchenko et al. 2009). NBN (founder) mutation, 657del5, is estimated to appear in 1 case per 177 newborn (Varon et al. 2000). Global MAF for the NBS causing variant is 0.04% (GnomAD v3.1.1) whereas in our results for the Polish population the frequency of this variant is almost seven fold higher at 0.27% (Fig. 10B). This is the first report of the allele frequency of the NBS causing variant for the Polish population, estimated on the basis of WGS.

Cystic Fibrosis (CF) is another example of an autosomal recessive disease, for which disease-causing allele spectrum was not derived from comprehensive genomic methods in Polish population. The analyses to date have utilized only targeted methods such as INNOLIPA CFTR19 test or Sanger Sequencing of chosen exons (NBS CF working group et al. 2013), without providing data on frequencies of intronic and regulatory site variants. Cystic fibrosis is the most frequent metabolic disease in Poland, with the incidence estimated at 1 per 4394 (NBS CF working group et al. 2013) to 1 per 5000 individuals (Farrell 2008). CF affects mostly lungs and leads to progressive breathing problems and respiratory failure. The disease is caused by mutations located in *CFTR* gene (188 kb, 27 exons), for which more than 2000 variants have been identified to date according to CFTR Mutation Database (data from 28 of June 2021 http://www.genet.sickkids.on.ca/StatisticsPage.html). In our study, we observed that the cumulative allele frequency for three pathogenic ClinVar variants found in POL cohort (rs78655421, rs113993960, rs77010898) in the *CFTR* gene is significantly lower than GnomAD NFE population. Specifically, in the case of the most common NM_000492.3(CFTR):c.1521_1523delCTT (p.Phe508del, rs113993960), we observed that the allele frequency in Polish population is slightly lower (0.95%) than in European NFE (1.43%). However, much lower than previously reported for Polish population, described as almost 3% carrier frequency according to (NBS CF working group et al. 2013; Morral et al. 1994). Surprisingly, in our study, Slavic specific NM_000492.3:c.54-5940_273+10250del (CFTRdele2,3) (Dörk et al. 2000) variant was not found. The reported mutation frequencies for the Polish population for p.Phe508del and CFTRdele2,3 are 57-62% and 1.8-6.2% respectively (Bobadilla et al. 2002; NBS CF working group et al. 2013). The very low frequency of CFTRdele2,3 deletion may be the reason why it was not detected in our cohort of 943 individuals.

### Relationship of the Polish and global populations

In order to explore the diversity of the Polish population and global relationships between POL and other world populations, we performed the principal component analysis (PCA) based on common variants (MAF>1%). In the first comparison with continental populations, we observed that the POL cohort is homogenous and clustered within the European population (Fig. 11A and 11B). After prediction using a random forest method only one sample was located in the AMR population cluster. In PCA of European subpopulations, almost all POL samples (938 out of 943) were clustered with other European ancestries, with 496 individuals belonging to the GBR, 427 to the CEU, 12 to the TSI, and 3 to the IBS subpopulations. Five samples were closer to non-European populations. The results are presented on Fig. 11 (C, D).

**Figure 11.**
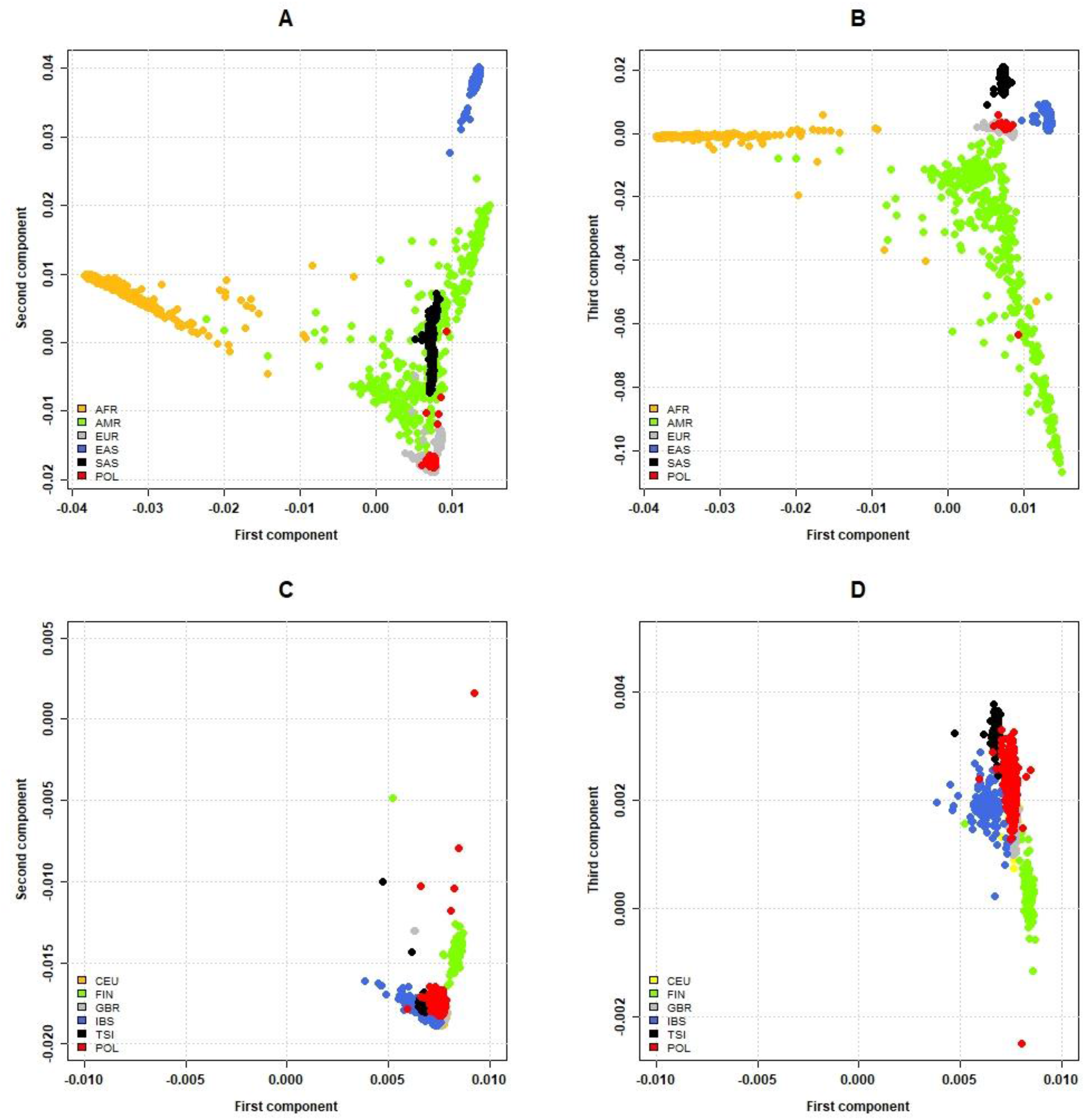
Clustering of samples among the World and European subpopulations based on the random forest and PCA prediction. The comparison of (A) the first and second principal components for continental populations; (B) the first and third principal components for continental populations; (C) the first and second principal components for the European subpopulations; (D) the first and third principal components for the European subpopulations.

The above-mentioned results were in accordance with the Fst statistics. Considering continental populations, POL was most similar to the European (an average Fst of 0.004), and least similar to the African and East Asian populations (average Fst 0.056 and 0.050 respectively). An average Fst statistics calculated over sliding windows of 1,000 SNPs is presented on Supplementary Figure S6. Furthermore, we compared the Polish cohort with other European subpopulations from the 1000 Genomes dataset and observed high average Fst between the cohorts (Table 2).

**Table 2:** Genomes project/world dataset.Fst statistics calculated for Polish cohort vs European subpopulations from the 1000.

| Population | CEU | FIN | GBR | IBS | TSI |
| --- | --- | --- | --- | --- | --- |
| Fst | 0.007 | 0.010 | 0.007 | 0.009 | 0.009 |

Additionally, density-based spatial clustering among POL individuals (Supplementary Fig. S7) was performed, confirming homogeneity of the cohort with the exception of a few outliers. The average Fst statistics between the POL cluster and the outliers was 0.012, suggesting a non-significant difference between the individuals and the rest of the cohort. The results of all three approaches (PCA, Fst and density-based spatial clustering) agree in demonstrating that POL is highly homogeneous and samples used in the analysis belong to the European populations, both in terms of continental and populational clustering.

We also performed admixture analysis in the unsupervised mode with the 943 unrelated individuals from POL and either the total 1000 Genomes dataset (Supplementary Fig. S8 .A) or the European populations (Supplementary Fig. S8.B). Compared against the world populations, the POL cohort was similar to the European populations at low K values (K = 2 to 5), and at K above 5 (favoured by the cross-validation analysis for the world dataset) forms a distinct cluster, with some common ancestry with GBR and CEU, and also FIN. In the analysis with only the European populations it formed a distinct cluster that was essentially homogeneous for the values of K less or equal 4. The lower K values were favoured by cross-validation results for the European dataset (the lowest CV error value was for K = 3).

## Discussion

Although large scale whole genome sequencing projects have arisen worldwide during the last decade, e.g ‘100 000 Genomes’ project in the UK, ‘1+ Million Genomes’ Initiative in Europe or ‘All of Us’ in the United States, country or region-oriented sequencing initiatives were not performed in many areas. Genome of the Netherlands (GoNL) project characterized the genomes of 769 Dutch ancestry individuals (The Genome of the Netherlands Consortium 2014) in 2014. In 2015 the genomes of 2 636 Icelanders were sequenced and described (Gudbjartsson et al. 2015), and in 2016 the population of Sardinia was characterized based on sequencing of 2 120 individuals (Sidore et al. 2015). These country based sequencing analyses significantly improved the SNP imputation performance, confirmed a high degree of geographic clustering of recent mutations and reliably discovered *denovo* mutations (Gudbjartsson et al. 2015; Sidore et al. 2015; The Genome of the Netherlands Consortium 2014), building local genomic resources for clinical genetics and fostering research that was not possible before.

In this study we present results of whole-genome sequencing and comprehensive analyses of a wide spectrum of genomic variation in 1079 individuals of Polish ancestry. We detected and genotyped small and structural variants (SV), runs of homozygosity, mitochondrial haplogroups, and Mendelian inconsistencies, among others. Allele frequencies for 943 unrelated individuals were released as a publicly available resource, The Thousand Polish Genomes (POL) database - the first effort of this scale completed for the Polish population. Two similar initiatives were launched in Poland in recent years, but their results are not openly available to the scientific community. The first one, the Polgenom^10^ project, ran in years 2013-2016, collected WGS data from 130 individuals of age 90+ and offered limited paid access to the data. In 2016 a large initiative entitled the Genomic Map of Poland^11^ was launched, aiming at sequencing a total of 5000 individuals, but its results are not available yet. In general, Central European populations are underrepresented in worldwide genomic databases such as GnomAD, HapMap or 1000 Genomes and the presented work fills this gap, and equips researchers with large and freely available reference dataset of variant frequencies.

The sequencing data gives a unique opportunity to interrogate rare variation. The rare alleles are considered relatively new in the population (Veltman and Brunner 2012) and they tend to cluster geographically more than the common alleles (Gravel et al. 2011; Mathieson and McVean 2012; The 1000 Genomes Project Consortium 2015). In the first phase of the POL project, which is described in the presented article, we focused on the identification of rare diseases-related variants, which may appear more frequently in Polish population in comparison to other European populations. We identified 736 pathogenic variants according to ClinVar, but more than 70% of them were present in a single person only, hindering reliable frequency estimates on a cohort of this size. The allele frequencies of the remaining pathogenic variants were up to 1.17 % (22/1886), which may have visible clinical consequences in the population. Among 16 pathogenic variants in the ACMG actionable genes identified in POL only 1 variant reached the frequency of 0.15 % (3/1886) (rs1217805587 in *BRCA1* gene). The overall incidence of secondary findings in Polish population matched the ACMG estimations. We suggest that national guidelines, available in Polish language, should be issued to address the importance of genetic counselling for clinical WES/WGS results concerning incidental findings specific to our population, as availability to whole genome analyses will successively increase.

The analysis of OMIM AR disease genes let us identify as many as 22 genes (Supplementary Table S5) with significantly different cumulative allele frequencies in POL vs European NFE populations. Variants of most of these genes are associated with the founder’s effect in the central european population and have shown the most striking differences between the POL population and the rest of Europe. The whole set of 22 genes may be taken into consideration by researchers and clinicians interested in specific rare diseases and following their incidence in the population.

Undoubtedly, apart of small variants, a valuable part of the released data is the structural variant (SV) dataset, called jointly on all unrelated samples in the cohort. Historically, SVs have been understudied due to methodological limitations. However, they collectively affect a larger part of the genome than the small variants (Sudmant et al. 2015), which we also observed in this cohort. Compared to other large-scale studies of SVs (Chiang et al. 2017; Collins et al. 2020; Hehir-Kwa et al. 2016), we report lower SV counts per individual, and larger fractions of affected genomic sequence, which may stem from methodological differences that are known to considerably affect structural variant detection (Cameron et al., 2019). We chose a single-tool approach with joint genotyping of the SVs, which could have resulted in higher false-positive rates and lower sensitivity than ensemble methods used in other studies (Collins et al., 2020; Sudmant et al., 2015). However, considering the uniqueness of this data and the role it could play, for instance for rare-variant filtering, we decided to release the unfiltered SV frequencies. In future releases of the database, the spurious SV calls will be marked, allowing researchers to more precisely compare SV characteristics between cohorts. We also note that further analyses combined with populational aCGH or optical mapping data are needed to explore and validate the diversity and clinical utility of SV patterns in Polish population and broader context as their pathogenic potential and involvement in complex genetic traits is still poorly understood.

It was previously described that the Polish population is relatively homogenous, with a low level of genetic diversity (Jarczak et al. 2019). Small genetic differences are present at regional level and are in correlation to patterns of migrations in the past (Jarczak et al. 2019). In the current study the analysis of mtDNA haplogroups was performed on 943 unrelated individuals. For the majority of people in our cohort (44 %) the haplogroup H was assigned, which is consistent with previous results for Polish and Slavic populations (Grzybowski et al. 2007; Jarczak et al. 2019; Mielnik-Sikorska et al. 2013). Other studies on mitochondrial DNA show that Poles as a population are characterized by different European haplogroups, with dominance of West Eurasian, Central and Eastern European haplogroups (Jarczak et al. 2019; B. A. Malyarchuk et al. 2002; Mielnik-Sikorska et al. 2013). It was also shown that Polish population is almost indistinguishable from other European nations, except for the sub-haplogroups U4a and HV3a which are predominantly found in Poles and Russians (B. Malyarchuk et al. 2008; B. A. Malyarchuk et al. 2002). In this project we confirmed only the presence of the U4a haplogroup (28 individuals), which is assumed to be of central-european origin (B. A. Malyarchuk et al. 2002).

The analyses of Polish population similarities to other populations performed in presented work clearly proved the homogeneity of POL as well as the closest ancestry with GBR and CEU populations. The admixture analysis also confirmed that the Polish cohort is closest to the other European populations in the 1000 genomes dataset. The Polish cohort forms a homogeneous cluster that is distinct from the other European populations, again confirming its low level of genetic differentiation. Due to the lack of other samples from Central and Eastern Europe in this dataset, making any detailed inference about the ancestry of the Polish population compared to the rest of the continent is impossible at the moment. In comparison to populations from northwestern and southern Europe that are present in the 1000 Genomes dataset, the Polish population is expected to have more traces of the steppe ancestry (Lazaridis, 2018), but the populations available in the dataset are not sufficient to test this hypothesis. One cannot also exclude the possibility that the distinct clustering of the Polish cohort is related to the differences in WGS data processing methods used to obtain different datasets.

Runs of homozygosity (ROH) analysis identifies the stretches of contiguous homozygous sites in an individual and are used to measure the level of inbreeding and recessive inheritance. The longer the total length of ROH in individuals the closer the inbreeding in a population, which increases the risk of recessive disease and decreases reproductive fitness of the offspring (Hoffman et al. 2014). The Polish population characterised in this study perfectly matches the European ROH characteristics). This suggests that the inbreeding level and potential risk of recessive diseases caused by homozygosity is relatively low. There is however some risk that sampling bias could have influenced the ROH analysis as most of the volunteering participants were city inhabitants. On the contrary ROH density increases in rural areas, less mobile and more prone to close-kin marriages. Unfortunately, this type of bias occurs in most large scale genomic studies to date (Dean et al. 2017).

As with many populational, volunteer-based projects, our data are not bias free. Our cohort was primarily recruited to perform the GWAS study on COVID-19 susceptibility, therefore we didn’t focus on collecting samples from Polish minorities like Kaschubians, Karaims, Lemkos or Polish Jews. We have also not corrected the much higher proportion of donors recruited from municipal areas compared to rural, as recruitment of volunteers from small villages is a cumbersome, costly process due to lack of trust and no means of effective project promotion in these communities. Also, the proportion of individuals from different voivodeships is skewed in favor of Masovia and Greater Poland, probably because of more intense promotion of the project that was conducted there. As a result, the far-eastern voivodeships were underrepresented which could have led to missing variants that could have been native e.g. to Tatar or Russian populations. Moreover, the cohort of more than 1000 genomes, however impressive, does not enable to reliably estimate the prevalence of variants less common than 0,5%, which could be very useful in clinical setting to re-evaluate and reclassify many VUS variants (variant of uncertain significance) to benign site of the spectrum.

Although in terms of sample size our project does not compare to the world’s largest, it remains one of the largest open allele-frequency datasets generated on high-coverage WGS data and the largest of Slavic population. Results presented in this paper are only the tip of an iceberg and should be taken as an invitation for further explorations. Our main goal was to generate an open-source dataset available for all researchers around the world, thus, filling in an existing gap in world’s datasets so far lacking Eastern-European samples. We believe that such collaborative efforts will be soon performed in many countries delivering excellent results for future research of many disciplines.

## Acknowledgements

Authors would like to thank the medical personnel of Central Clinical Hospital of Ministry of the Interior and Administration in Warsaw, Multidisciplinary Municipal Hospital of Józef Struś in Poznań and ALAB Laboratoria Sp.z o.o. as well as SiePomaga Foundation for collaboration and blood sample collection.

This work was partially supported by the Polish National Science Centre grant No. SZPITALE-JEDNOIMIENNE/2/2020 and by the Medical Research Agency grant No 2020/ABM /COVID19/0022

## Footnotes

1 https://github.com/MNMdiagnostics/1000PolishGenomes

2 https://www.bioinformatics.babraham.ac.uk/projects/fastqc/

3 https://github.com/brentp/smoove

4 https://github.com/brentp/smoove

5 https://github.com/MNMdiagnostics/1000PolishGenomes

6 https://www.omim.org/, accession date 26-04-2021

7 https://panelapp.genomicsengland.co.uk/panels/, accession date 27-06-2021

8 https://github.com/MNMdiagnostics/1000PolishGenomes

9 https://www.omim.org/, accession date 26-04-2021

10 https://polgenom.pl

11 https://www.genompolski.pl

