## Supplementary Material for "‘The Thousand Polish Genomes Project’ - a national database of Polish variant allele frequencies"

### Supplement

The supplement contains 10 figures and 4 tables.

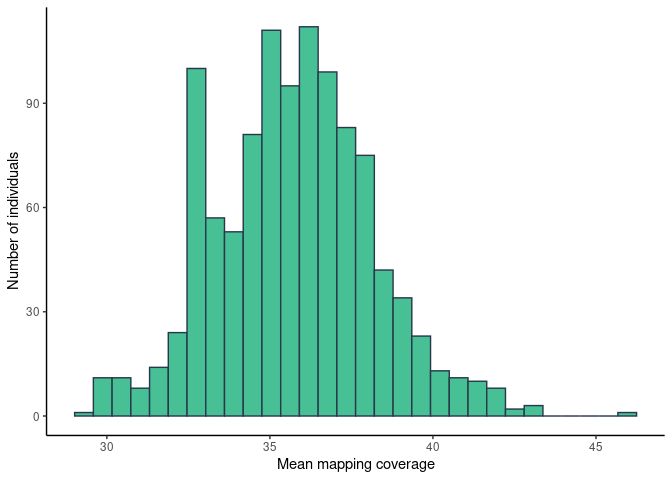

**Fig.S1:  Distribution of mean mapping coverage in the analysed cohort.**

**Table S1: Coverage statistics for all 1079 samples analysed in the project.**

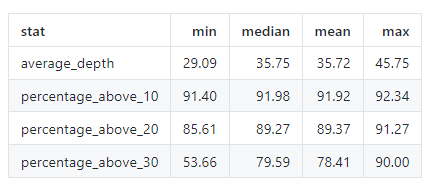

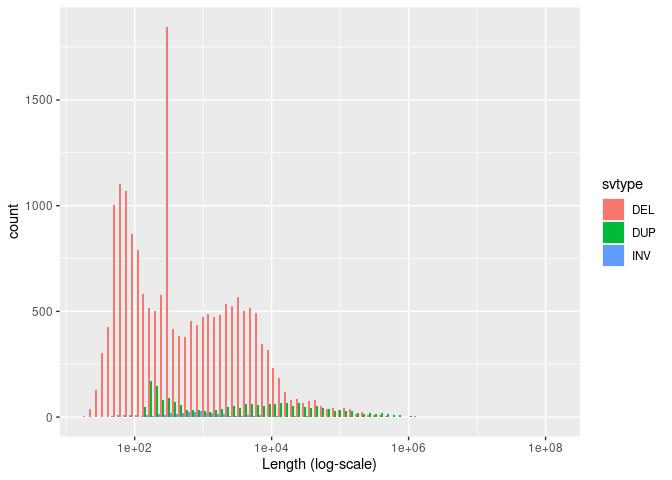

**Fig. S2.A: Distribution of structural variant (SV) lengths (log10 scale) for three SV types: deletions (DEL), duplications (DUP), and inversions (INV) among the analysed cohort.**

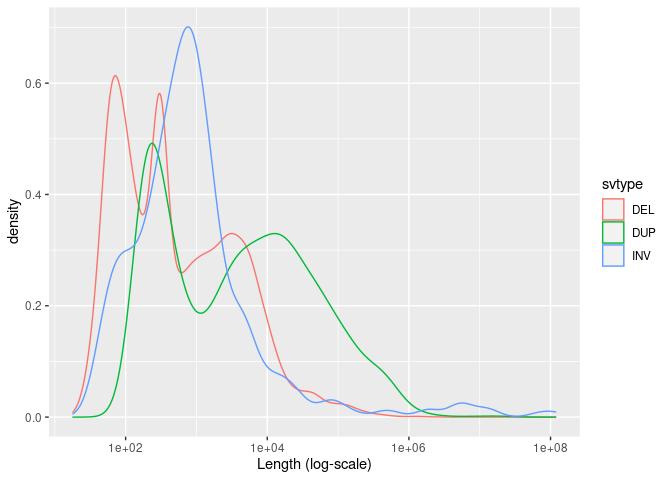

**Fig. S2B: Distribution of structural variant (SV) lengths (log10 scale) presented as density for the three SV types: deletions (DEL), duplications (DUP), and inversions (INV) among the analysed cohort.**

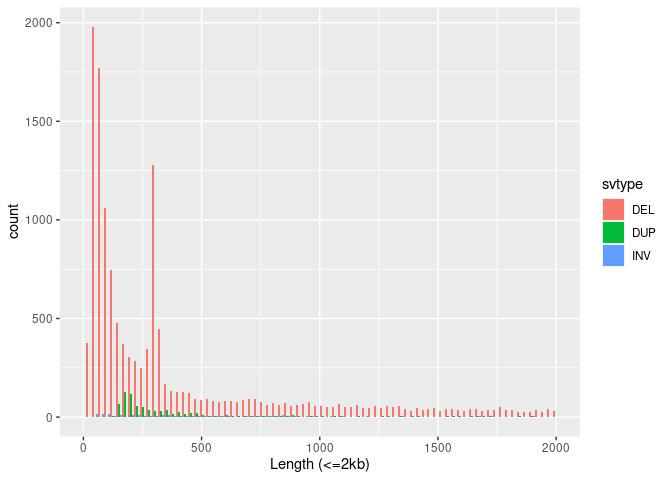

**Fig. S2C:** **Distribution of structural variant (SV) lengths in the range 0-2000bp, for the three SV types: deletions (DEL), duplications (DUP), and inversions (INV) among the analysed cohort.**

**Supplementary table S2:** **SNV counts per individual.**

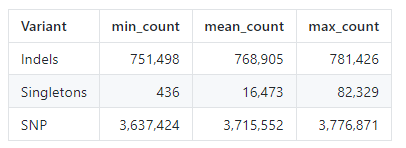

**Supplementary Table S3: Total number (count), number of high quality (count.PASS) and percentage of high quality (pct.pass) structural variants in the dataset. BND-breakend, DEL-deletion, DUP-duplication, INV-inversion.**

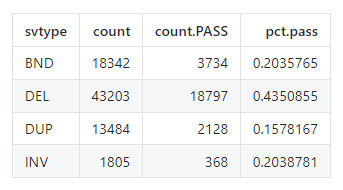

| **Supplementary Table S4: Numbers of structural variants per individual. BND-breakend, DEL-deletion, DUP-duplication, INV-inversion, het-heterozygous genotype, hom-homozygous genotype.**  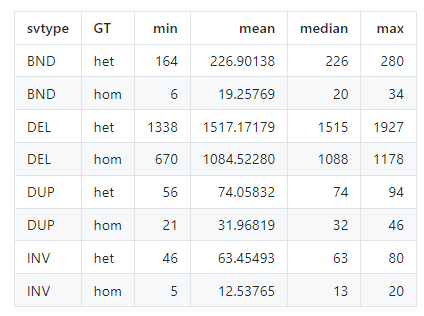 |
| --- |

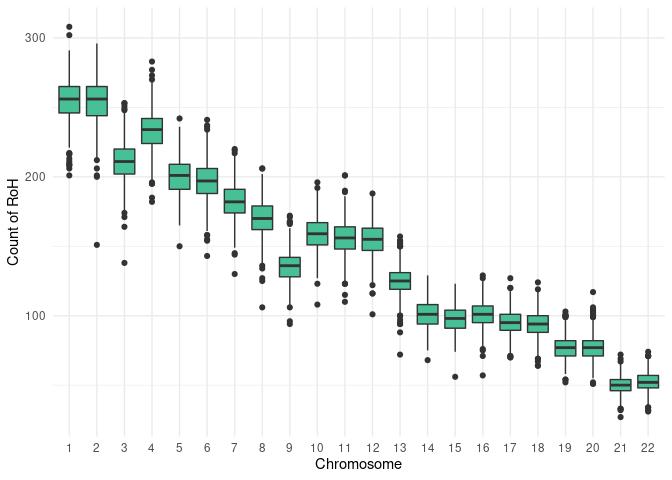

**Fig. S3: Number of Runs of homozygosity (RoHs) per chromosome. RoH analysis identifies the stretches of contiguous homozygous sites in an individual and are used to measure the level of inbreeding and recessive inheritance.**

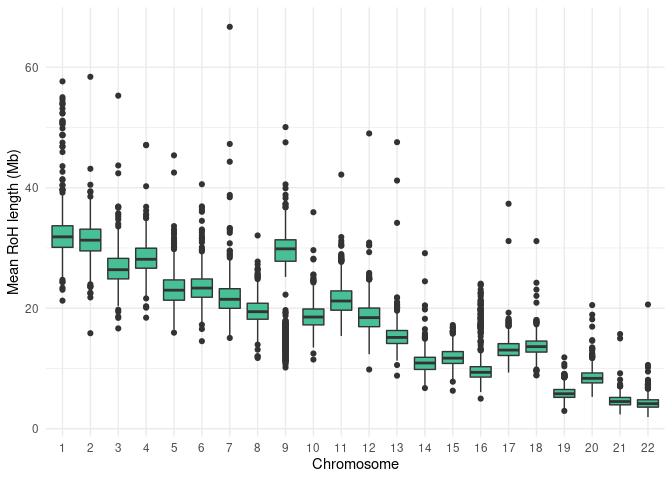

**Fig. S4: Average size of Runs of homozygosity (RoHs) per chromosome**.

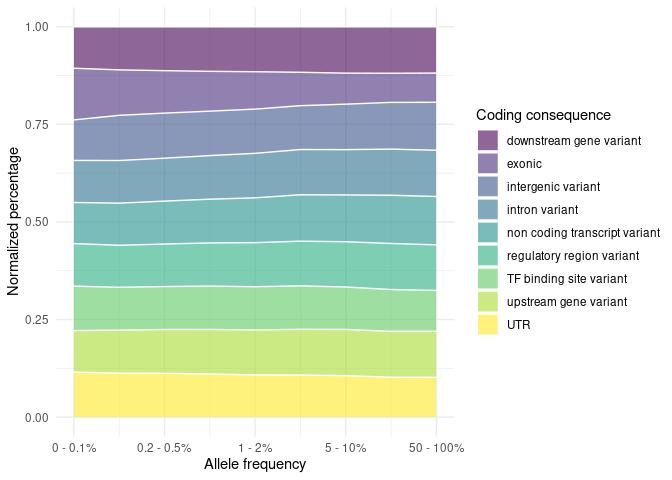

**Fig. S5: Distribution of variant consequences across allele frequency spectrum.**

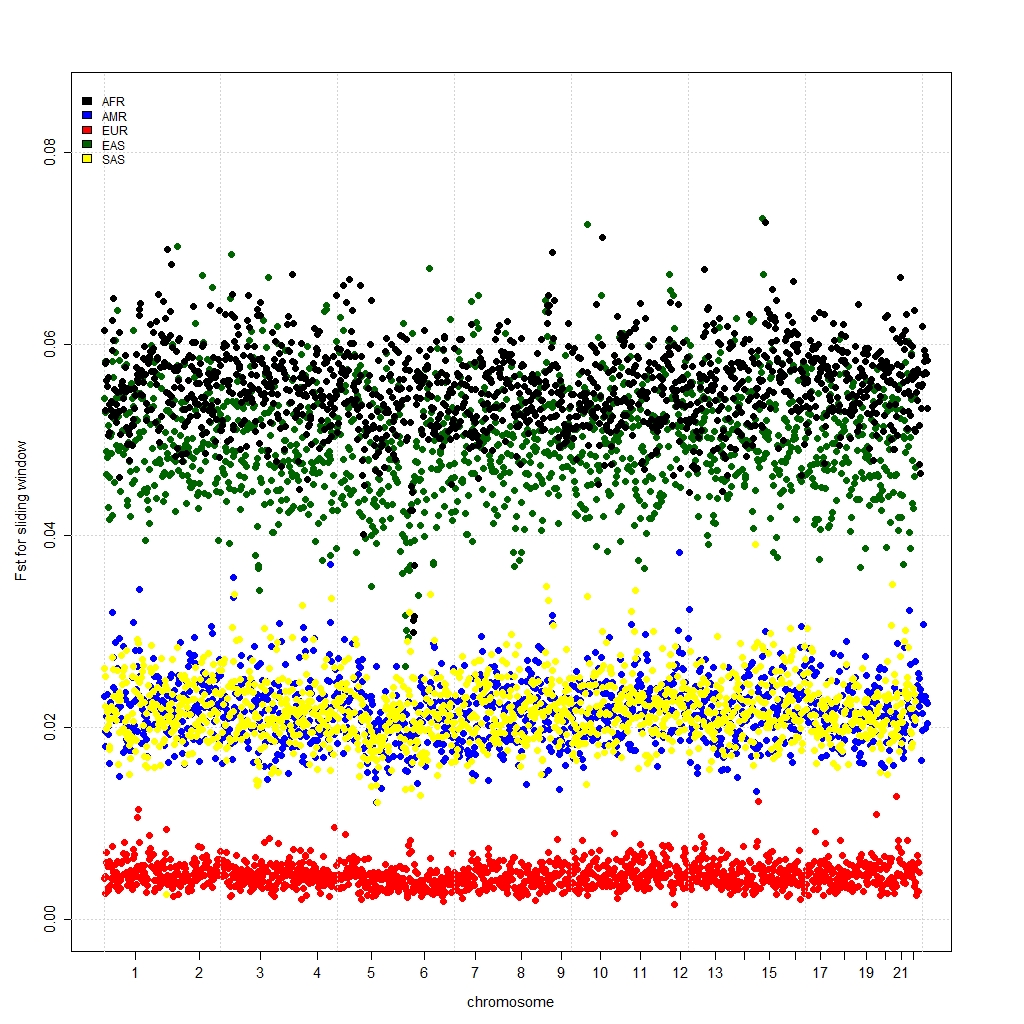

**Fig. S6: An average Fst statistics calculated over sliding windows of 1,000 SNPs for POL vs continental populations.**

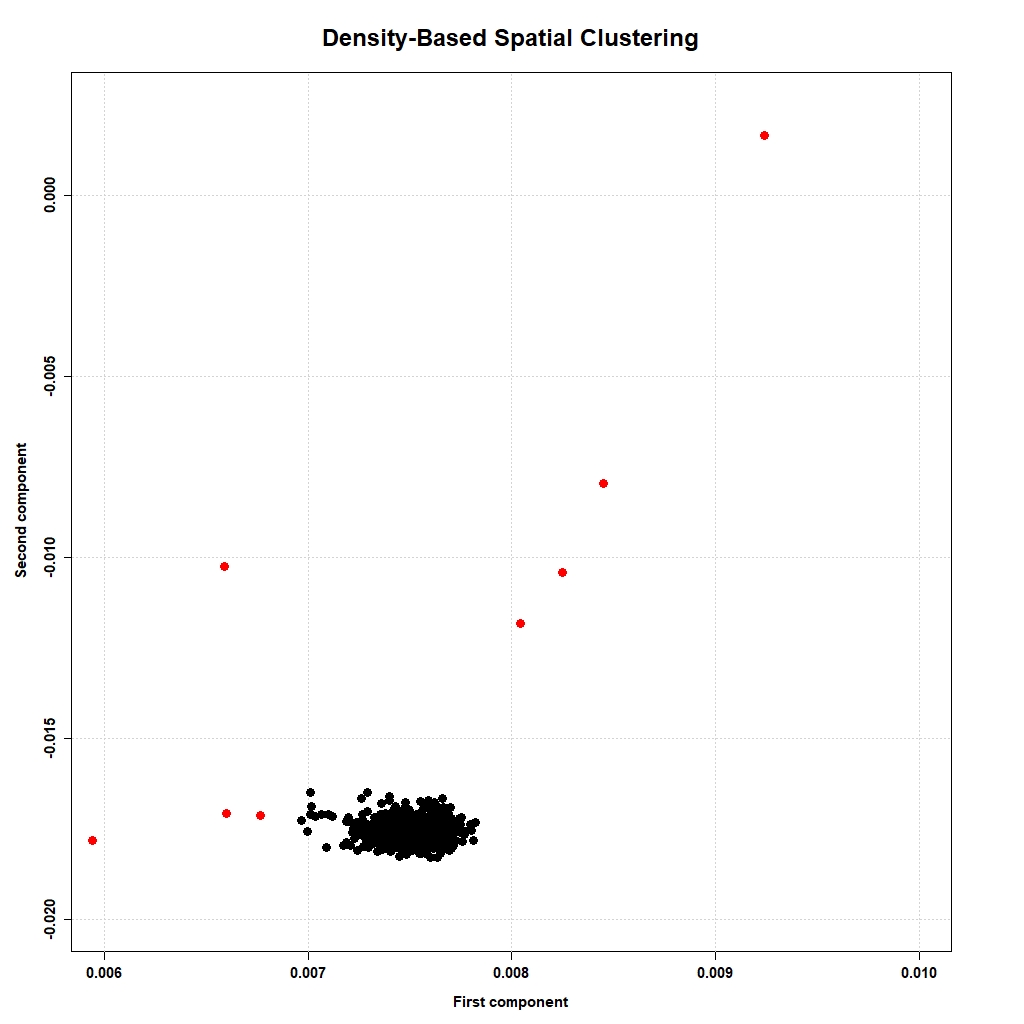

**Fig. S7: Density-based spatial clustering of POL cohort.**

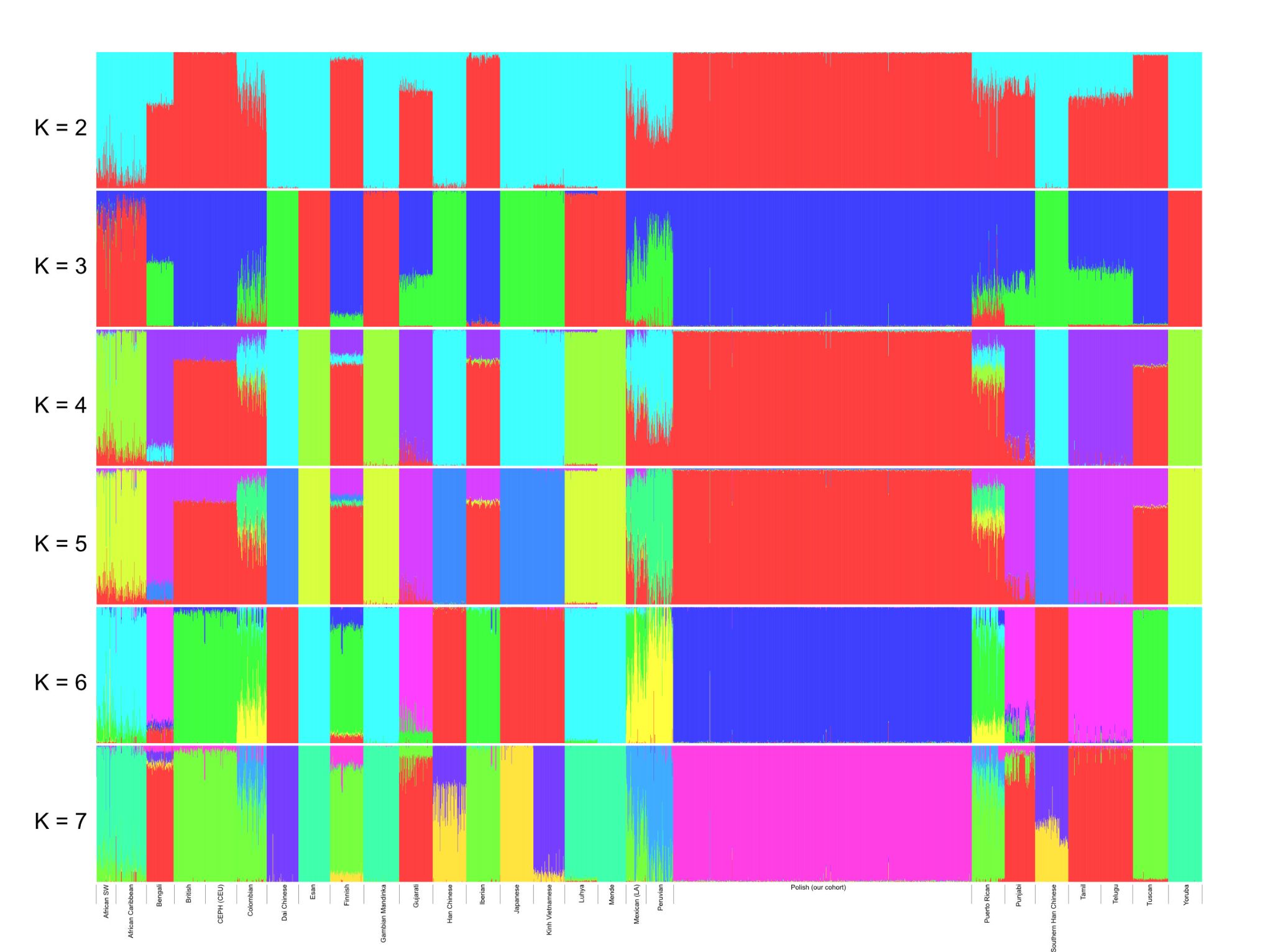

**Fig. S8 A: Admixture (Alexander et al., 2009) plots of the Polish cohort together with the 1000 Genomes world dataset. K >= 7 minimizes the cross validation error value.**

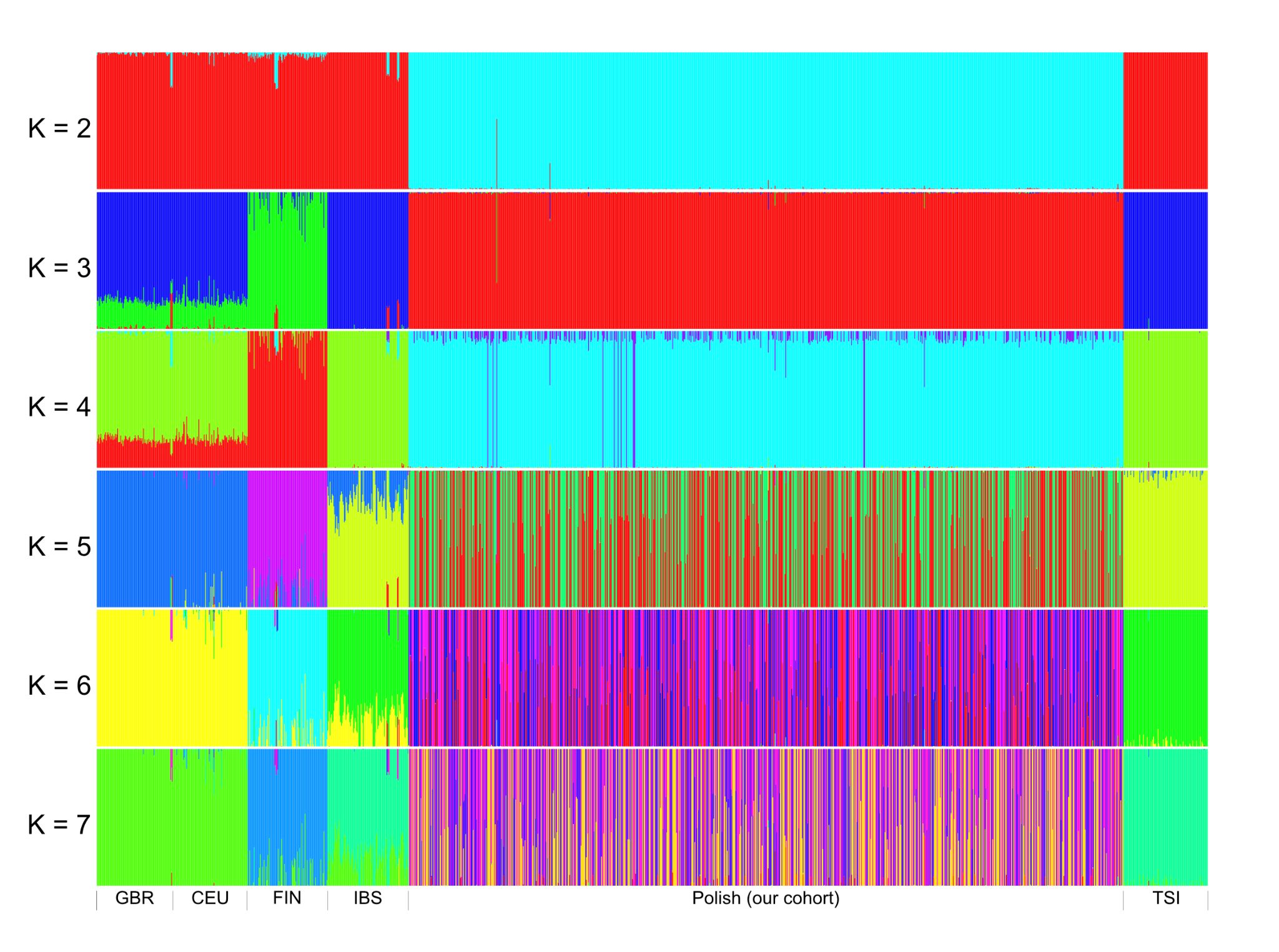

**Fig. S8 B: Admixture (Alexander et al., 2009) plots of the Polish cohort together with the European populations from the 1000 Genomes world dataset. K = 3 minimizes the cross validation error value.**
